# Microglia regulate neuronal activity via structural remodeling of astrocytes

**DOI:** 10.1101/2025.02.18.638874

**Authors:** Ning Gu, Olena Makashova, Celeste Laporte, Chris Q. Chen, Banruo Li, Pierre-Marie Chevillard, Graham Lean, Jieyi Yang, Calvin Wong, Jonathan Fan, Behrang Sharif, Susana Puche Saud, Misha Hubacek, Katrina Y. Choe, Arkady Khoutorsky, Charles W. Bourque, Masha Prager-Khoutorsky

## Abstract

Neuron-glia interactions play a central role in regulating synaptic transmission and neuronal excitability. Structural plasticity of astrocytes is associated with numerous physiological and pathological conditions, however, the mechanism underlying this process remains unknown. To examine the basis for structural astrocyte plasticity, we used the classic example of the loss of astrocytic processes that takes place in the hypothalamic magnocellular system during chronic high-salt intake. We discovered that a high-salt diet triggers a local accumulation of reactive microglia around vasopressin-secreting neurons, but not in other brain areas. Microglia phagocytose astrocytic processes, reducing astrocytic coverage of vasopressin neurons. The pruning of astrocytic processes impairs synaptic glutamate clearance, enabling activation of extrasynaptic glutamate NMDA receptors and increasing the activity of vasopressin neurons. Inhibiting microglia-mediated astrocyte pruning attenuates the increased neuronal activity and vasopressin-dependent hypertensive phenotype of rats fed high-salt diet. Thus, microglia orchestrate neuron-glia interactions and regulate neuronal activity through astrocyte pruning.

## Introduction

Neuron-glia interactions are crucial for regulating neuronal activity to control brain function in physiological and pathological conditions^1,2^. For example, astrocytes play a central role in modulating synaptic transmission, neurotransmitter clearance, and synaptic plasticity^3^. Therefore, the structural remodeling of astrocytes, which occurs under various conditions, can strongly influence neural activity^4^.

One of the classical systems where structural plasticity of astrocytes and its functional outcomes have been investigated is the magnocellular hypothalamo-neurohypophysial system comprising supraoptic (SON) and paraventricular (PVN) nuclei, harboring magnocellular neurons releasing vasopressin (VP) or oxytocin^5^. A striking feature of the astrocytes surrounding magnocellular neurons in the SON and PVN is their ability to undergo robust remodeling in response to the animal’s physiological state demanding large and sustained increases in hormonal release^6,7–9^. This plasticity is characterized by the reversible withdrawal of astrocytic processes that envelop magnocellular neuron somata and dendrites, resulting in an increased number of direct soma-to-soma appositions, as well as a higher number of synapses^8^. Previous studies have demonstrated a significant reduction in astrocytic coverage of oxytocin neurons during lactation and chronic dehydration, indicating that this structural plasticity facilitates and/or makes hormonal release more efficient^1,10–14^. It has also been described in dehydration caused by water deprivation, supporting sustained increases in VP secretion to minimize water loss when an animal has limited access to water, as well as following exposure to a high-salt diet (HSD)^8,15,16^. Although this remarkable plasticity was first documented nearly five decades ago^16–19^, the mechanism underlying this process remains unknown, and our understanding of its role is still incomplete.

Here, we addressed these questions by examining astrocyte structural plasticity in response to HSD. We found that astrocyte remodeling is mediated by reactive microglia, which phagocytose astrocytic processes surrounding magnocellular VP neurons. This leads to impaired astrocytic glutamate clearance and enhanced activation of extrasynaptic NMDA glutamate receptors, thus contributing to increased activation of VP neurons and VP-dependent elevation in blood pressure. Inhibiting microglia-mediated pruning of astrocytes attenuates changes in VP neuron activity and HSD-induced hypertension in rats, uncovering a new mechanism by which microglia orchestrate structural plasticity and neuron-glia interactions.

## Results

### High-salt intake induces loss of astrocyte processes

To analyze astrocyte structural plasticity in the magnocellular system we exposed rats to HSD for 7 days by adding 2% NaCl to their drinking water. Consistent with previous ultrastructural studies^7,8,16–21^, immunohistochemical examination of the magnocellular system in rats fed HSD revealed a substantial decrease in astrocytic coverage, as visualized by glial fibrillary acidic protein (GFAP) in the SON (Fig. 1A) and PVN (Fig. S1A and S1B). To confirm this observation, we evaluated additional astrocytic markers, including S100 calcium-binding protein β (S100β), aldehyde dehydrogenase 1 family member L1 (ALDH1L1), glutamate transporter 1 (GLT1), and vimentin (Fig. 1B-E). Our data show reduced immunolabeling of these astrocytic proteins in the SON of rats fed HSD (Fig. 1F-J). The HSD-mediated decrease in astrocytic coverage was further validated by Western blot analysis, which showed reduced levels of the astrocytic proteins GFAP, GLT1, and ALDH1L1 in the SON of rats fed HSD compared to control diet (Fig. S1C and S1D). A significant decrease in astrocytic coverage is observed as early as 5 days but not at 2 days after exposing rats to HSD (Fig. S2A and S2B). Notably, the total number and density of astrocyte cell bodies in the SON and PVN remain unchanged after HSD (Figs. S2C and S2D), suggesting that HSD does not cause the elimination of astrocytes but rather triggers a reduction in astrocytic processes surrounding magnocellular VP neurons.

**Fig. 1:**
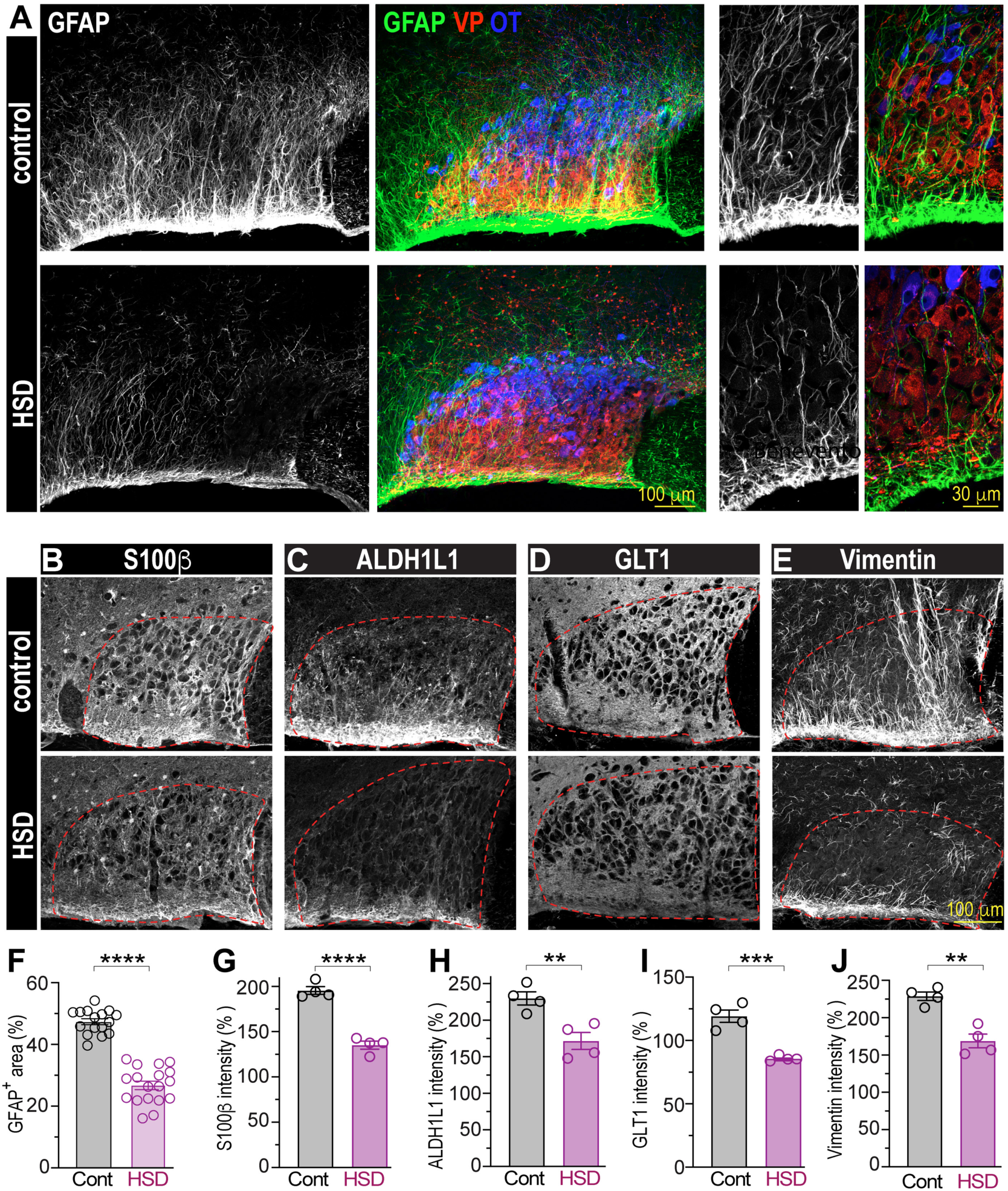
Rats fed high-salt diet exhibit decreased astrocyte coverage. (**A**) Coronal rat brain sections immunolabeled to visualize astrocytes (GFAP, white or green) and neurons secreting VP (red) or oxytocin (blue) in the SON of control rats (top panels) or rats fed HSD (7 days of 2% NaCl in the drinking solution, bottom panels). Additional astrocytic markers S100β (**B**), ALDH1L1 (**C**), GLT1 (**D**), and vimentin (**E**) visualized in the SON of rats fed control diet (upper panels) and HSD (bottom panels). Red dotted lines outline VP neuron location. (**F**) Quantification of GFAP in the SON of rats fed control diet or HSD (n=16 control, n=18 HSD rats). (**G**-**J**) Quantifications of S100β, ALDH1L1, GLT1, and vimentin in the SON of rats fed control diet or HSD, n=4 rats/group. Data presented as mean ± SEM, analyzed by Student’s t-test, \*\*\*\**P* < 0.0001, \*\*\**P* < 0.001, \*\**P* < 0.01, each data point represents one rat. See also Figs. S1 and S2.

### High-salt intake causes accumulation of reactive microglia around VP

Previous studies postulated that decreased astrocytic coverage is mediated by astrocyte retraction^5,7,16,18,19,22^, however evidence supporting this notion has been lacking. Microglia are the main immune cell type in the central nervous system (CNS) and have the capacity to engulf and phagocytose cellular elements^23,24^. Therefore, we hypothesized that microglia engulf astrocytic processes in HSD. To test this hypothesis, we examined the effect of HSD on microglia organization in the SON and PVN of rats. We found that Iba1-positive microglia accumulate in the SON and PVN of HSD-fed animals but not in the surrounding areas (Fig. 2A). Since Iba1 is expressed not only by microglia but also by macrophages, we used both Iba1 and the pan-microglial marker TMEM119 to confirm that HSD increases the density of microglia around VP neurons (Fig. 2B-G). In addition to the increased density of microglia around VP neurons in HSD (Fig. 2A and 2E-G, and S3A-C), HSD induced changes in microglial morphology from a highly ramified shape with multiple thin branches to a less complex morphology with shorter and thicker processes (Fig. 2D), indicative of their activation^25^. Sholl analysis was used to quantify these changes in microglial morphology, showing a significant decrease in the number of Sholl intersections in microglia from HSD-treated rats (Fig. 2H). These changes in microglial morphology appeared gradually and were detectable as early as 2 days after exposing rats to HSD, with microglia progressively acquiring a more reactive morphology by 7 days of HSD (Fig. S3D). Notably, while reactive microglia accumulated around VP neurons in the SON and PVN, their density and morphology remained unchanged outside these nuclei, where microglia remained in a “resting/naïve” state (Fig. S4). These results demonstrate that feeding rats HSD triggers the local accumulation of reactive microglia in brain nuclei containing magnocellular VP neurons.

**Fig. 2:**
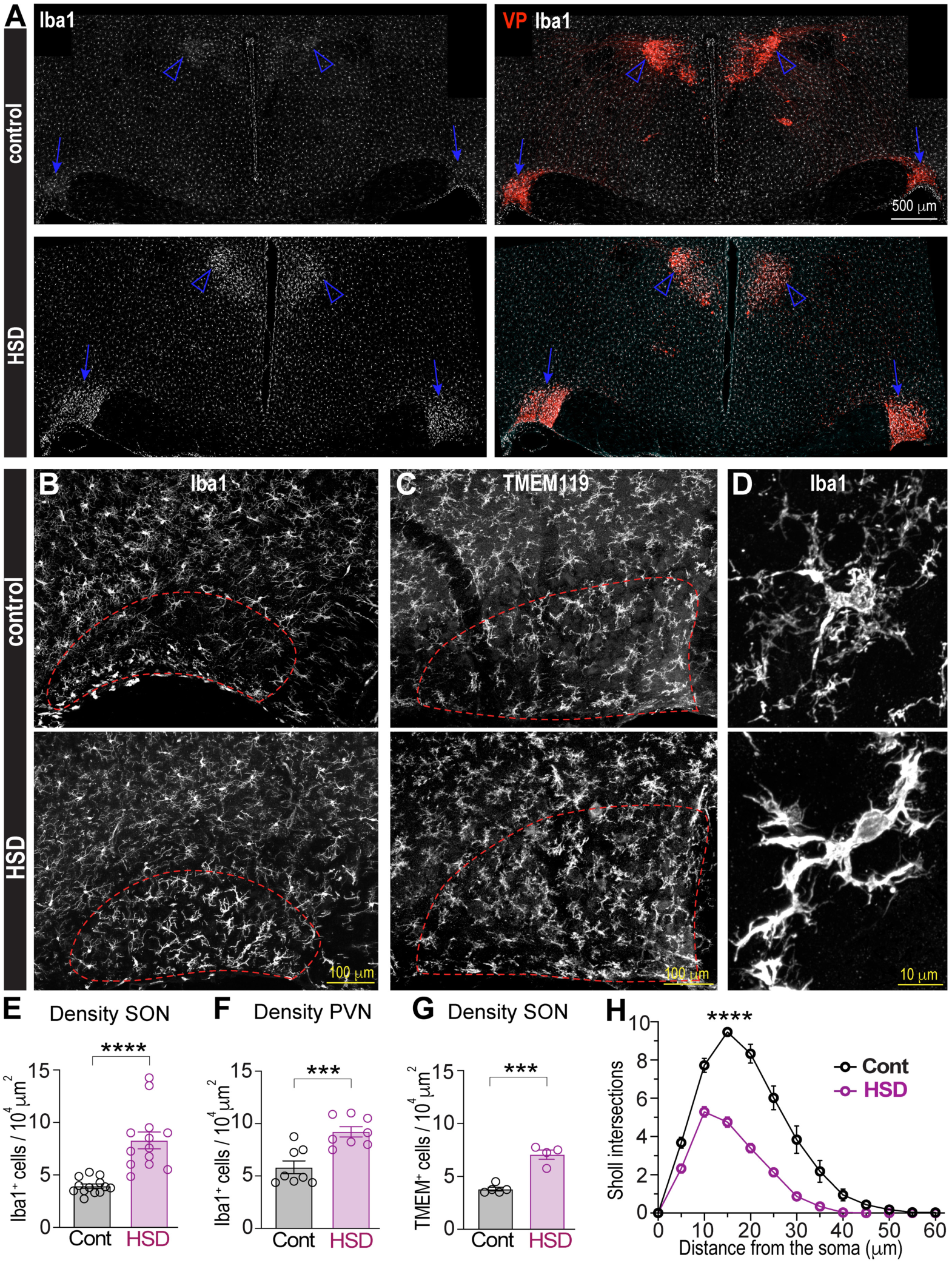
Local accumulation of reactive microglia in the SON and PVN of rats fed high-salt diet. (**A**) Low-magnification images of hypothalamic brain sections showing Iba1-positive microglia (white) and VP neurons (red) in the SONs (arrows) and PVNs (arrowheads). Rats fed control diet (upper panels) show a homogeneous distribution of microglia, while in HSD (bottom panels), microglia accumulate around VP neurons of the SON and PVN. (**B**-**C**) Confocal images of SON sections from the control (upper panels) and HSD-treated (bottom panels) rats show Iba1 (**B**) and TMEM119 positive microglia (**C**), red dotted line outlines VP neuron location. (**D**) High magnification Airyscan images illustrate HSD-induced changes in Iba1-positive microglia morphology from highly ramified in control (upper panel) to less complex shape with shorter and thicker processes, typical of reactive microglia in the SON from HSD rats (bottom panel). (**E**-**G**) Quantification of Iba1-positive microglia density in the SON (**E**, n=13 rats/group), PVN (**F**, n=8 rats/group), and TMEM119-positive microglia density in the SON (**G**, n=4 rats/group) in control or HSD rats, analyzed by Student’s t-test. (**H**) Distribution of Sholl intersection number by the distance from microglia soma in the SONs from control and HSD rats (n=5 rats/group, 20 cells per animal), p < 0.0001 by two-way ANOVA with Šídák’s multiple comparisons post-hoc test). Data presented as mean ± SEM, \*\*\**P* < 0.001, \*\*\*\**P* < 0.0001, each data point represents one rat. See also Figs. S3 and S4.

### Microglia inhibition attenuates the loss of high-salt diet-induced astrocyte processes

To study whether microglia play a role in the loss of astrocytic processes around magnocellular VP neurons, we inhibited microglia using pharmacological agents and examined if this prevents the loss of astrocytic processes induced by HSD. To inhibit microglia, we used PLX3397, an inhibitor of the colony-stimulating factor 1 receptor (CSF1R), a key regulator of microglia proliferation, activation, and survival^26,27^. The rats were administered PLX3397 (30 mg/kg/day, intraperitoneally, i.p.) or the vehicle daily for 3 days, then exposed to either control diet or HSD for an additional 7 days, with PLX3397 administration continuing throughout the dietary protocols (for a total of 10 days). Treatment of rats fed HSD with PLX3397, but not the vehicle, prevented the accumulation of reactive microglia and blocked the loss of astrocytic processes around VP neurons (Fig. S5).

To further confirm these observations, we used an additional CSF1R inhibitor, PLX5622, which is less effective overall but more specific in targeting CSF1R^28^, with a 10-fold lower affinity for related kinases compared to PLX3397^29^, and is therefore considered more efficient in depleting microglia^28,29^. The rats were administered with either the vehicle or PLX5622 (twice a day, for a total of 100 mg/kg/day, i.p.) for 3 days, after which PLX5622 or vehicle injections continued during exposure to HSD or control diet. Treatment of HSD-fed rats with PLX5622, but not the vehicle, not only prevented the accumulation of reactive microglia but decreased their density compared to control (Fig. S6A-C). Similar to PLX3397, treatment of HSD-fed rats with PLX5622 also inhibited the loss of astrocytic processes around VP neurons (Fig. S6A and S6D).

Microglia are the only cell type expressing CSF1R in the CNS^30,31^; however, systemic administration of CSF1R inhibitors can also affect peripheral macrophages^32^. Thus, to avoid effects on peripheral macrophages and selectively target microglia in the CNS, we administered an additional CSF1R inhibitor, BLZ945^33^, directly into the brain. To this end, rats were implanted with an intracerebroventricular (i.c.v.) cannula in the lateral ventricle. Two weeks after recovery, they received daily i.c.v. injections of BLZ945 (0.3 mg/kg/day) or vehicle for 3 days. On the fourth day, rats were exposed to either control diet or HSD for an additional 7 days, while daily BLZ945 administration continued throughout the dietary protocols. Treatment of HSD-fed rats with BLZ945, but not the vehicle, prevented the accumulation of microglia in the SON (Fig. 3A-C) and PVN (Fig. S7A-C). BLZ945 treatment completely reversed HSD-induced changes in microglia morphology, suggesting that it prevented microglia activation in response to HSD (Fig. 3D). Strikingly, this inhibition of microglia blocked the loss of the astrocytic marker GFAP around VP neurons of the SON (Fig. 3A and 3E) and PVN (Fig. S7A and S7D) of rats exposed to HSD. The effect of microglial inhibition on HSD-induced decreases in GFAP coverage was detected as early as 5 days after the initiation of the dietary protocol (Fig. S8). Furthermore, in addition to GFAP, microglia inhibition with BLZ prevented the HSD-induced loss of all examined astrocytic markers in the SON, including S100β, ALDH1L1, GLT1, and vimentin (Fig. S9A-D). These data suggest that HSD-induced accumulation of reactive microglia mediates the reduction in astrocytic coverage around VP neurons in the SON and PVN.

**Fig. 3:**
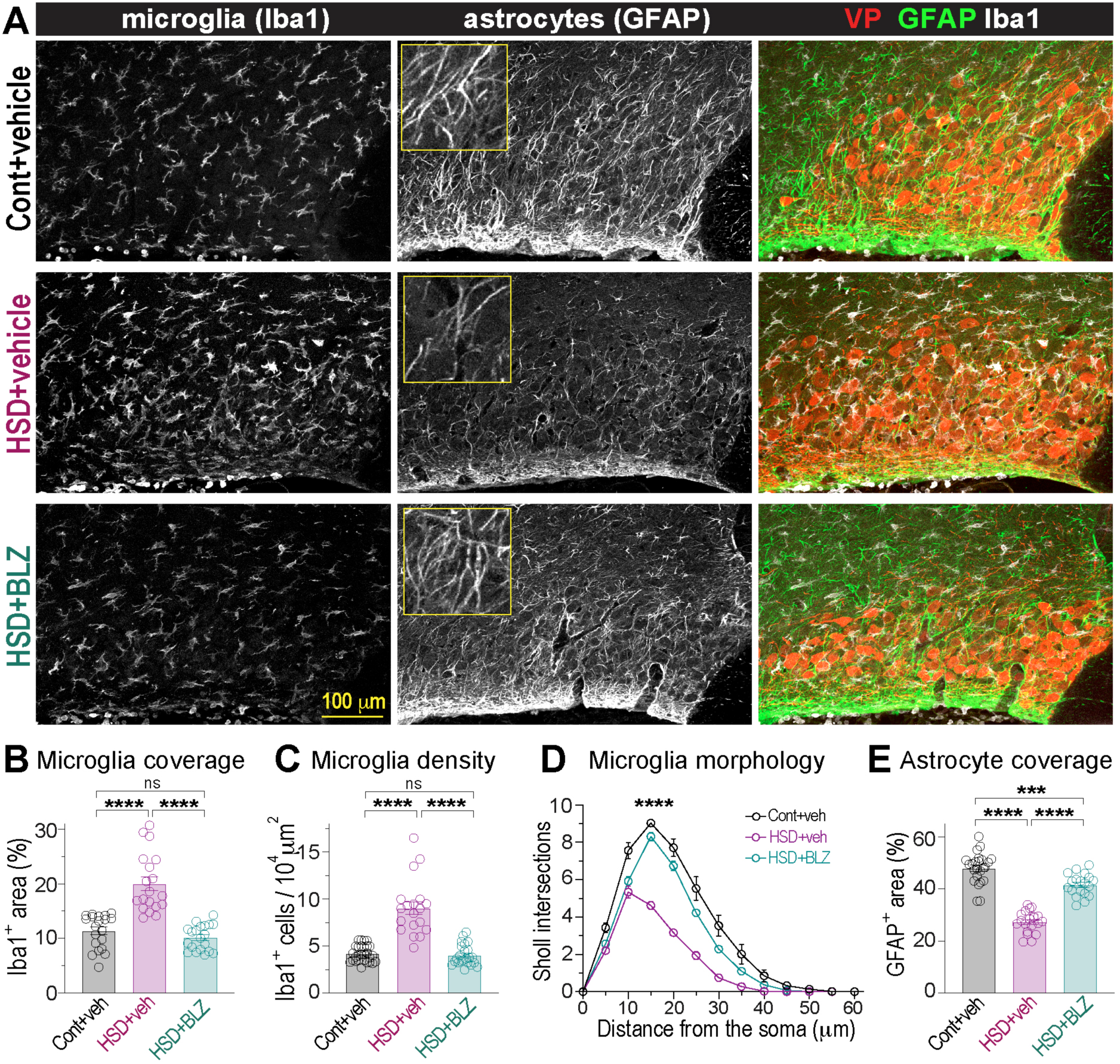
Microglia inhibition blocks astrocyte remodeling in high-salt diet. (**A**) SON sections from control (cont) and HSD-fed rats treated with vehicle i.c.v. (veh), and HSD rats treated with 0.3 mg/kg/day BLZ945 i.c.v. (HSD+BLZ), labeled with markers of microglia (Iba1), astrocytes (GFAP), and VP neurons. Quantification of microglia coverage, analyzed as Iba1-immunopositive SON area (**B**), and microglia density (**C**) in rats exposed to different treatments (n=19 rats/group, one-way ANOVA with Tukey’s post-hoc test). (**D**) Distribution of Sholl intersection number by the distance from microglia soma in the SONs from vehicle-treated control and HSD-fed rats and BLZ-treated HSD rats (n=5 rats/group, 20 cells per animal), p < 0.0001 by two-way ANOVA with Tukey’s post-hoc test. (**E**) Astrocyte coverage analyzed as GFAP-immunopositive SON area in rats exposed to different treatments (n=19 rats/group, one-way ANOVA with Tukey’s post-hoc test). Data presented as mean ± SEM, \*\*\**P* < 0.001, \*\*\*\**P* < 0.0001, ns, not significant; each data point represents one rat. See also Figs. S5, S6, S7, S8, and S9.

### Microglia phagocytose astrocytic processes during high-salt intake

Microglia phagocytose neuronal elements under various physiological conditions (e.g. developmental synaptic pruning and plasticity)^23^, as well as in disease states (e.g. multiple sclerosis, Alzheimer’s disease)^24,34,35^. Accordingly, the enhanced phagocytic capacity of reactive microglia is associated with an increased number of CD68-positive (CD68+) phagocytic lysosomes within microglia in the phagocytic state. Assessing CD68+ lysosomes has therefore been used to quantify the phagocytic capacity of microglia^36,37^.

To determine whether HSD stimulates microglia to become phagocytic and engulf astrocytic processes surrounding VP neurons, we analyzed whether microglia in this condition contain CD68+ lysosomes. Super-resolution imaging with Airyscan combined with 3D reconstruction of individual microglia using IMARIS revealed that the number of CD68+ lysosomes was higher in SON microglia from rats fed HSD compared to those on control diet (Fig. 4A-C, Video <u>S1</u> and <u>S2</u>). Moreover, HSD significantly increased the volume occupied by phagocytic lysosomes within the microglial cytoplasm (Fig. 4D). These HSD-induced increases in the number and volume of CD68+ lysosomes were abolished when rats were administered BLZ945 i.c.v. but not the vehicle (Fig. 4A-C, Video <u>S3</u>). Strikingly, 3D colocalization analyses showed that most CD68+ lysosomes within SON microglia from HSD-fed rats contained astrocytic markers GFAP and S100β (Fig. 4A, 4E, and 4F, and Video <u>S2</u>). We found very little or no astrocytic components in microglia from rats fed control diet. Additionally, BLZ945 inhibited HSD-induced increases in astrocytic material within microglial CD68+ lysosomes (Fig. 4A, 4E, and 4F). To examine whether microglia in HSD-treated rats also engulf neuronal components, we analyzed if CD68+ lysosomes colocalize with the general pan-neuronal marker NeuN and VP, which is abundantly present in the dendrites of magnocellular VP neurons. Importantly, while most microglial CD68+ lysosomes contained significant amounts of astrocytic markers GFAP and S100β in HSD (Fig. 4E and F, and Fig. S10A), CD68+ lysosomes showed minimal co-localization with NeuN and VP in SON microglia in both control and HSD-fed rats (Fig. S10B and S10C). These results indicate that HSD stimulates microglial phagocytic activity, leading to the engulfment of astrocytic components and microglia-mediated pruning of astrocyte processes.

**Fig. 4:**
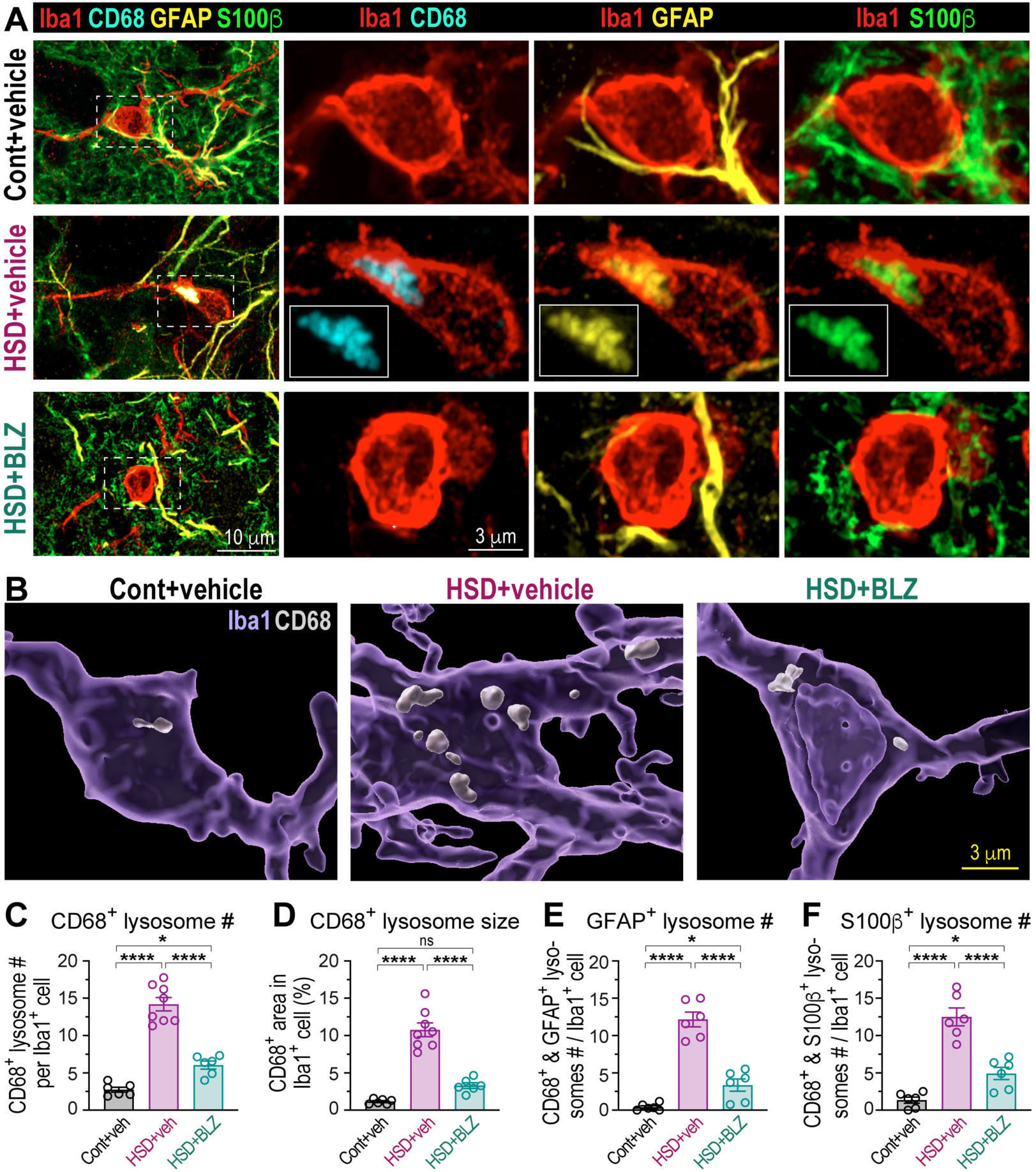
Microglia phagocytose astrocytic processes in high-salt diet. (**A**) Super-resolution Airyscan images show microglia (Iba1, red), microglia-specific lysosomal marker CD68 (cyan), and astrocyte markers GFAP (yellow), and S100β (green) in the SONs from control (cont) and HSD-fed rats treated with vehicle i.c.v. (veh), and HSD rats treated with 0.3 mg/kg/day BLZ945 i.c.v. (HSD+BLZ). (**B**) IMARIS 3D surface reconstruction of microglia and CD68+ lysosomes from control and HSD-fed rats treated with BLZ or vehicle showing Iba1 (translucent purple), CD68 (opaque white); see also 3D reconstructions Video S1, S2, and S3. Bar graphs show the number of CD68+ lysosomes per microglia (**C**), the proportion of microglial area occupied by CD68+ lysosomes (**D**), and the number of CD68+ lysosomes colocalizing with GFAP (**E**) and S100b (**F**) in above conditions, n=6 rats/group, one-way ANOVA with Tukey’s post-hoc test. Data presented as mean ± SEM, \*\*\*\**P* < 0.0001, \**P* < 0.05, ns, not significant; each data point represents one rat. See also Fig. S10 and Videos S1, S2, and S3.

### Microglia-mediated pruning of astrocytic processes impairs synaptic glutamate clearance

Chronic hypertonic stimuli, such as HSD, enhance the osmosensitivity of magnocellular VP neurons^38^ and increase their action potential firing *in vivo*^39^. Whether changes in astrocyte coverage can contribute to this increased neuronal activity in HSD remains unclear. Previous studies showed that the degree of astrocytic coverage can govern the availability of transmitters in the extrasynaptic space, contributing to the regulation of synaptic efficacy and neuronal activation^40,41^. Our data show that the loss of astrocyte processes in HSD is manifested by decreased levels of several astrocytic proteins, including GLT1 (Fig. 1I and Fig. S1D). Since GLT1 is the primary astrocytic glutamate transporter responsible for glutamate clearance in the synapse^42^, we hypothesized that in HSD, microglia-mediated astrocytic pruning impairs synaptic glutamate clearance by GLT1. As a result, spillover glutamate can activate extrasynaptic NMDA receptors (NMDARs) on VP neurons and induce a non-inactivating depolarizing current^43,44^ that contributes to the excitation of these neurons.

To test whether HSD-induced astrocyte pruning impairs the ability of GLT1 to regulate VP neuron activity, we performed whole-cell current-clamp recordings from VP neurons in hypothalamic slices. VP neurons were identified by their expression of eGFP^45^. Consistent with previous reports, pharmacological inhibition of GLT1 with dihydrokainate (DHK), a specific inhibitor of GLT1, depolarized and increased the action potential firing rate of VP neurons in acute hypothalamic slices prepared from rats fed control diet (Figs. 5A and S11A). However, inhibiting GLT1 had no effect on VP neuron membrane potential and firing rate in slices prepared from rats fed HSD (Figs. 5B and S11B). These results indicate that astrocytic GLT1 restricts the activity of VP neurons in the control condition, but this regulatory mechanism is lost in HSD. Notably, blocking microglia-mediated pruning of astrocytes by i.c.v. administration of BLZ945 throughout 7 days of HSD completely restored the effect of GLT1 inhibition by DHK on VP neuron excitability, while treating rats fed control diet with BLZ945 had no effect on GLT1-mediated excitation of VP neurons (Figs. 5C-F, S11C-F). These data demonstrate that microglia-mediated pruning of astrocytes in HSD hinders GLT1’s capacity to prevent glutamate spillover. This impaired glutamate clearance by GLT1 results in spillover into the extrasynaptic space, which may activate extrasynaptic NMDARs.

**Fig. 5:**
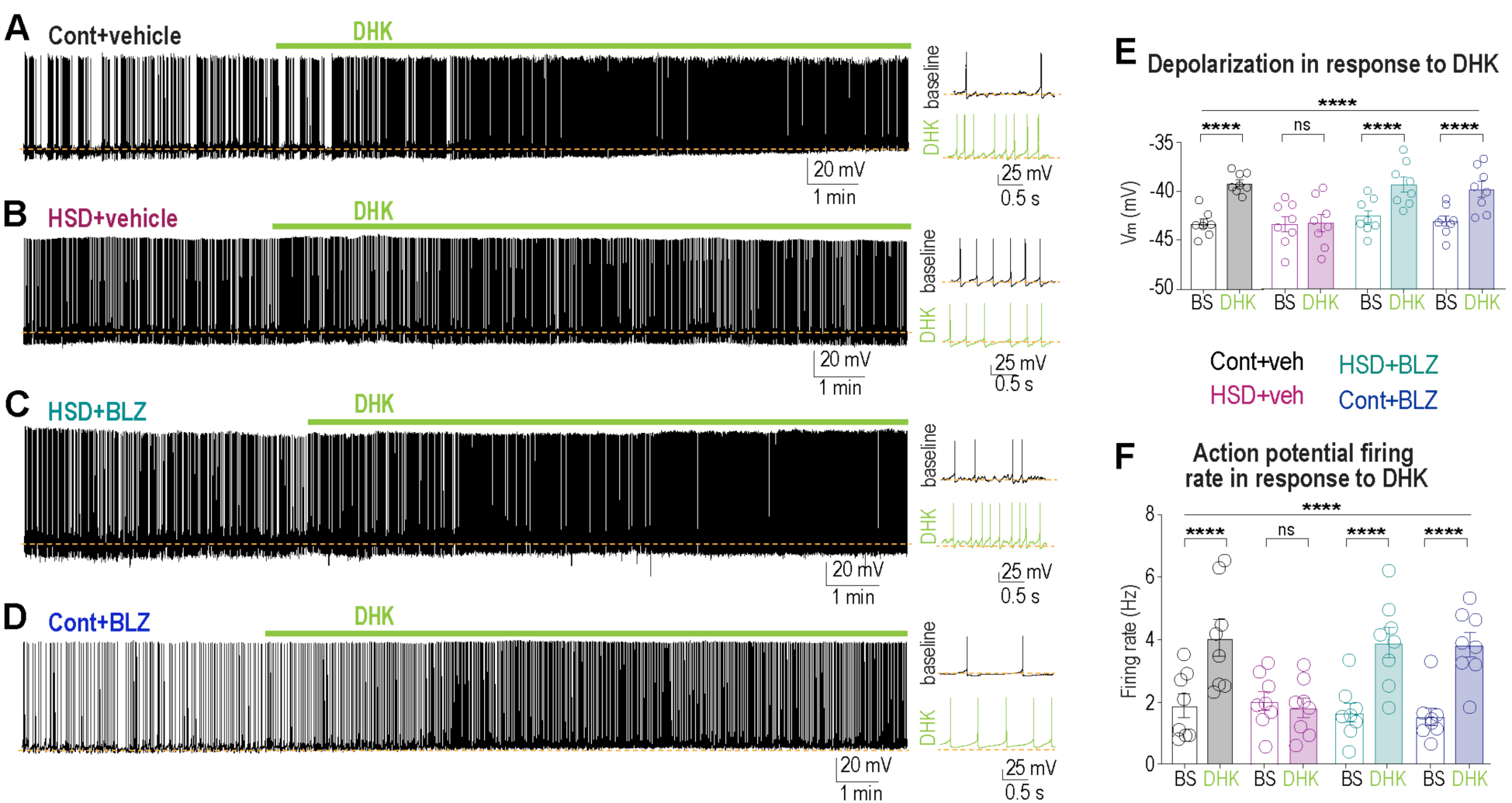
Microglia-mediated astrocyte pruning impairs synaptic glutamate clearance by astrocytic glutamate transporter. Left: Sample current-clamp traces show the effect of bath application of glial glutamate transporter blocker DHK (100 μM, green bar) on action potential firing of VP neurons in SON brain slices prepared from vehicle-treated control (**A**) and HSD (**B**) rats and BLZ945-treated HSD (**C**) and control (**D**) rats. Right: expanded 2-second sample traces before (black) and after (green) DHK application. Dashed lines show membrane potential at baseline. Bar graphs show changes in membrane potential (**E**) and firing rate (**F**) induced by DHK in SON VP neurons from vehicle-treated control and HSD and BLZ945-treated HSD and control rats (n=8 rats/group, repeated measures two-way repeated measures ANOVA with Šídák’s multiple comparisons post-hoc test). Data presented as mean ± SEM, \*\*\*\**P* < 0.0001, ns, not significant; each data point represents one rat. See also Fig. S11.

### High-salt diet-induced impairment of synaptic glutamate clearance increases activation of extrasynaptic NMDARs

To examine whether impaired glutamate clearance contributes to an enhanced activation of extrasynaptic NMDARs in HSD, we measured persistent NMDAR-mediated currents in VP neurons from control and HSD rats. To evaluate the contribution of tonic extrasynaptic NMDARs, we used ifenprodil, a selective blocker of NR2B NMDAR subunit^46^, which is a predominant subunit comprising extrasynaptic NMDARs^47,48^. To assess the contribution of different glutamatergic currents, VP neurons from SON slices were voltage-clamped to +40 mV. The baseline holding current was recorded, along with changes in the holding current in response to the sequential bath application of drugs targeting different subsets of glutamate receptors. First, extrasynaptic NMDA currents were selectively blocked with ifenprodil. The remaining (synaptic) NMDA currents were then inhibited with AP-5. Finally, all residual non-NMDA glutamate receptors were blocked by applying the general blocker kynurenic acid (Fig. 6A). Our results show that extrasynaptic NMDAR currents are significantly increased in VP neurons from HSD-fed rats compared to those on control diet, and this increase was fully reversed by blocking microglia-mediated pruning of astrocytes with BLZ945 (Fig. 6A-G). Also, we found that neither AP-5 sensitive currents, corresponding to synaptic ifenprodil-insensitive NMDARs, nor kynurenic acid, corresponding to non-NMDA (AMPA and kainate) currents, were altered in different treatments (Fig. 6F-G). These data demonstrate that HSD leads to an increased availability of extrasynaptic NMDARs, and preventing microglia-mediated pruning of astrocytic processes in HSD reverses this increase.

**Fig. 6:**
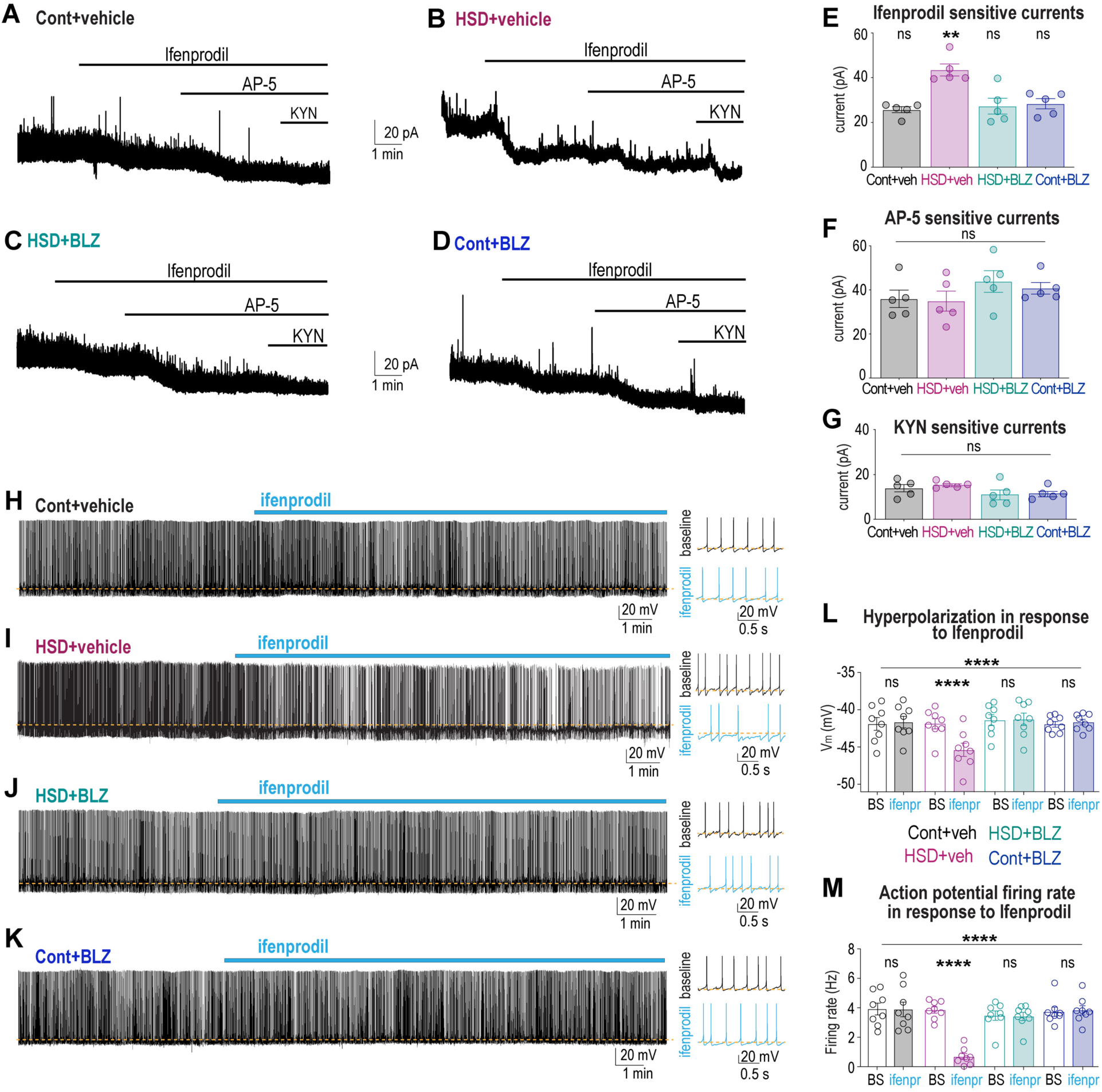
Microglia-mediated astrocyte pruning increases tonic excitatory extrasynaptic NMDAR currents in VP neurons. (**A-D**) Sample traces showing the tonic NMDA current (I_hold_) by sequential applications of ifenprodil (10 µM), AP-5 (100 µM), and kynurenic acid (1 mM) from voltage-clamped VP neurons (V_hold_ +40 mV) in SON brain slices obtained from vehicle-treated control (**A**) and HSD rat (**B**) and BLZ945-treated HSD (**C**) and control (**D**) rats. Bar graphs show changes in I_hold_ in response to the application of ifenprodil (**E**), AP-5 (**F**), and kynurenic acid (**G**), in VP neurons from vehicle or BLZ945-treated control and HSD rats (5 rats per condition, one-way repeated measures ANOVA with Tukey’s post-hoc test). **(H-K**) Left: Sample current-clamp traces show the effect of ifenprodil (10 μM, blue bar) on action potential firing of VP neurons in SON slices from vehicle-treated control (**H**) and HSD (**I**) rats and BLZ945-treated HSD (**J**) and control (**K**) rats. Right: expanded 2-second sample traces before (black) and after (blue) ifenprodil application. Dashed lines show membrane potential at baseline. Bar graphs show changes in membrane potential (**L**) and firing rate (**M**) induced by ifenprodil in VP neurons in SON brain slices from vehicle or BLZ945-treated control and HSD rats (n=8 rats/group, repeated measures two-way ANOVA with Šídák’s multiple comparisons post-hoc test). Data presented as mean ± SEM, \*\*\*\**P* < 0.0001, \*\**P* < 0.01, ns, not significant; each data point represents one rat. See also Fig. S12.

To test whether the elevated tonic activation of extrasynaptic NMDARs contributes to increased excitability of VP neurons in rats fed HSD, we examined whether VP neuron activity is affected by blocking extrasynaptic NMDAR currents with ifenprodil. Current-clamp recordings from identified VP neurons showed that inhibiting extrasynaptic NMDAR currents has no effect on the activity of VP neurons from rats fed control diet (Fig. 6H, L, M, and Fig. S12A). However, blocking extrasynaptic NMDARs in VP neurons from HSD-fed animals hyperpolarized and reduced action potential firing rate of these neurons (Fig. 6I, L, M, and Fig. S12B), revealing the presence of tonic excitatory currents in this condition. Blocking microglia-mediated pruning of astrocytes in HSD with BLZ945 eliminated ifenprodil-sensitive inhibition, while treating rats fed control diet with BLZ945 had no effect (Figs. 6J-M and S12C-F). These data indicate that microglia-mediated pruning of astrocytic processes impairs synaptic glutamate clearance, leading to the tonic activation of extrasynaptic glutamate receptors on magnocellular VP neurons, and contributing to the elevated firing activity of VP neurons in HSD.

### Microglia-mediated astrocyte pruning contributes to blood pressure increases in HSD

HSD has been shown to increase the firing rate of VP neurons *in vivo*, contributing to elevated blood pressure and hypertension^39^. To study whether microglia-mediated increases in VP neuron excitation contribute to elevated blood pressure in rats fed HSD, animals were implanted with blood pressure telemetry devices and subjected to either HSD or control diet (Fig. 7A and 7B). Consistent with previous studies, rats fed HSD for 7 days displayed elevated mean arterial pressure (MAP) (Fig. 7C and 7D). Treating rats with microglia inhibitors PLX3397 (i.p.) or BLZ945 (i.c.v.), but not vehicles, significantly attenuated increases in blood pressure induced by HSD (PLX3397: Fig. 7C, BLZ945: Fig. 7D). These findings show that microglia-mediated increases in the tonic activation of VP neurons contribute to elevated blood pressure in rats fed HSD.

**Fig. 7:**
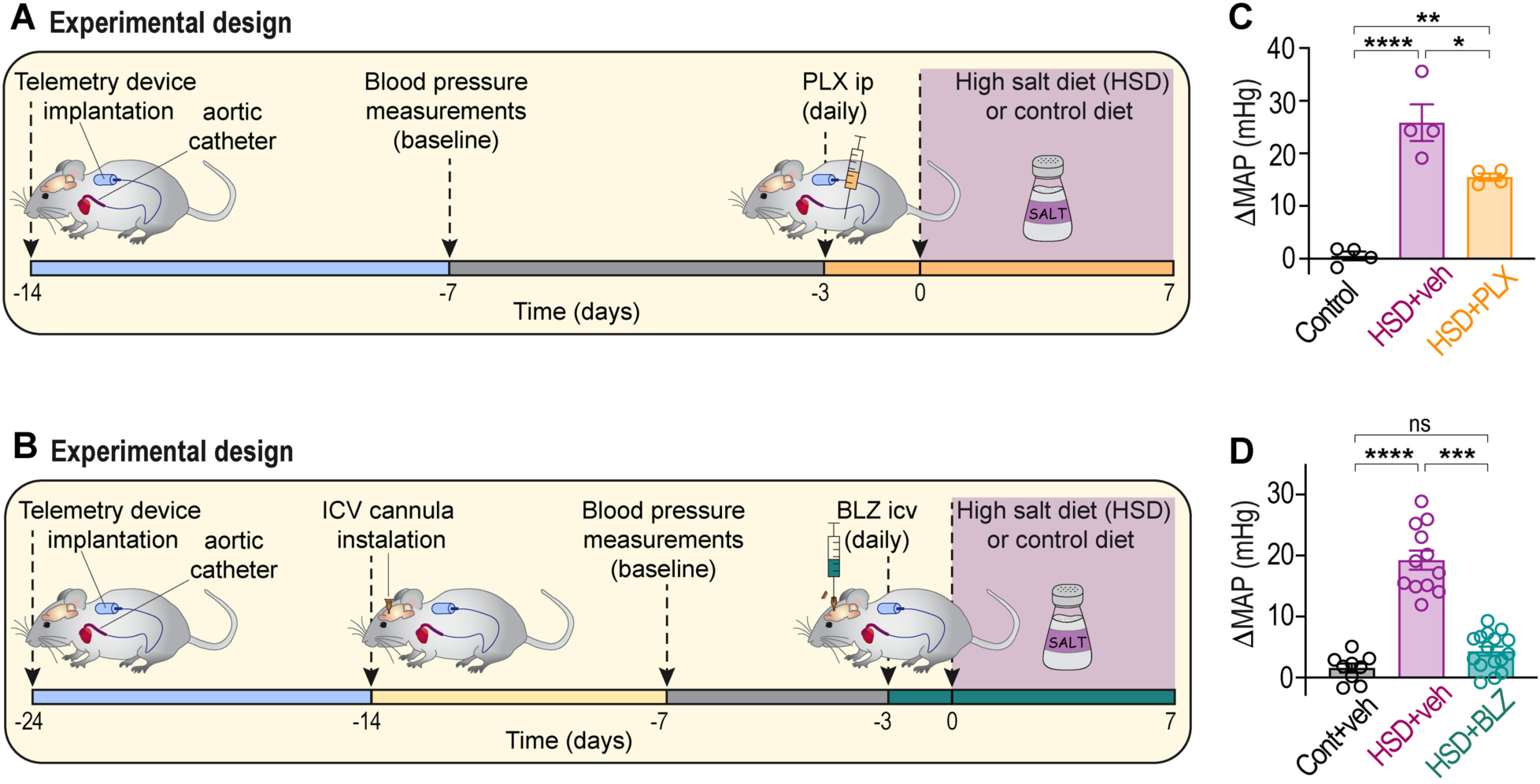
Blocking microglia-mediated astrocyte pruning prevents high-salt diet-induced increases in blood pressure. Schematic illustrations of the experimental design for blood pressure measurements in rats fed HSD and administered with PLX3397 i.p (**A**) and BLZ945 i.c.v (**B**). (**C**) Bar graphs show changes in the mean arterial pressure (MAP) in rats fed control diet or HSD and treated with 30 mg/kg/day PLX3397 or vehicle i.p. daily (n=4 rats/group). (**D**) Bar graphs show changes in the MAP in rats fed control diet or HSD and treated with 0.3 mg/kg/day BLZ945 or vehicle i.c.v. daily (n=9 rats for control, n=12 rats for HSD rats treated with vehicle, and n=16 HSD rats treated with BLZ945), analyzed by one-way ANOVA with Tukey’s post-hoc test. Data presented as mean ± SEM, each data point represents one rat, \*\*\*\**P* < 0.0001, \*\*\**P* < 0.001, \*\**P* < 0.01, \**P* < 0.05, ns, not significant.

## Discussion

Our study reveals that chronic high-salt intake, which is increasingly present in modern diets, is associated with a local neuroinflammatory process characterized by an accumulation of reactive microglia around magnocellular VP neurons in the SON and PVN. The microglia engulf and prune local astrocytic processes, impairing their ability to clear glutamate from the synaptic space. This ultimately causes glutamate to activate extrasynaptic NMDA receptors, contributing to the tonic activation of VP neurons. Furthermore, our findings show that this mechanism contributes to elevated blood pressure in rats fed HSD and may therefore be an important causal factor in salt-dependent hypertension.

### The central role of microglia in remodeling astrocyte network in HSD

Microglia participate in many critical brain functions, including neurogenesis, synaptogenesis, and myelination, through surveillance, release of soluble factors, and their capacity for phagocytosis during early postnatal development, aging, and disease conditions^49–52^. During development, microglia regulate the abundance of neuronal progenitors by clearing apoptotic or dying newborn neurons, removing non-functional synapses and remodeling synaptic circuits^36,53,54^, and phagocytosing developing oligodendrocytes^55^. In the adult CNS, microglia survey the environment, acting as first responders to acute injury and playing a central role in neurological diseases by clearing dead neurons, debris, and large extracellular aggregates via phagocytosis. These responses typically involve microglia transforming to a reactive state, retracting their processes, forming new motile protrusions, and becoming less ramified and more ameboid^49,50,56^. Previous studies have demonstrated that astrocyte-microglia communication is important during CNS development, and signaling from astrocytes to microglia promotes synapse engulfment by microglia during circuit maturation^57^. However, it remains unknown whether microglia can engulf astrocytic material during physiological or pathological plasticity.

Using a classical model where the astrocytic network undergoes substantial structural remodeling, we tested whether microglia can also phagocytose astrocytic material. Our study reveals that hypothalamic microglia can phagocytose and prune astrocytic processes in the adult rat brain. We found that subjecting rats to HSD leads to changes in microglia density and morphology, and these changes are specifically localized to areas occupied by VP neuron somata and dendrites in the SON and PVN (Fig. 2). These changes in microglia density and morphology are gradual, emerging after 5 days of HSD and peaking at day 7 (Fig. S3). Strikingly, microglia located in the adjacent areas, immediately outside the direct proximity of VP neurons, appear to be unaffected, as neither their morphology nor density changes even after 7 days of HSD (Fig. S4). Thus, while signals driving changes in microglia remain unknown, it is likely that these signals originate from the neighboring VP neurons. It is conceivable, that since magnocellular VP neurons are intrinsically osmosensitive^58,59^, they become more active in response to HSD-induced increases in blood osmolality^38^, secrete molecules that chemoattract and activate microglia, inducing their transition to a reactive phagocytic state and accumulation around VP neurons.

Our findings suggest that even at rest, SON microglia contain a low number of CD68+ lysosomes, which are much smaller than those found in reactive microglia after 7 days of HSD (Fig. 4 and Video <u>S1</u>). Some of these small lysosomes contain astrocytic material, suggesting that these microglia might be involved in constitutive astrocytic network remodeling under baseline, non-stimulated conditions. We found that some lysosomes rarely contained neuronal components, although to a much lesser extent than astrocytic material. Strikingly, we found that upon stimulation by HSD, there was a large increase in the total number and size of phagocytic lysosomes, and most of them contained astrocytic material (Fig. 4 and Video <u>S2</u>). These results are consistent with the previous observation reported that while HSD induces reduction in astrocytic coverage, it almost triples the number of synapses and significantly increases the size of neuronal somata and dendrites (neuronal hypertrophy)^8,10,19–21,60^. Hence, this raises the question of how these reactive microglia can specifically target local astrocytes while avoiding engulfing dense neuronal elements within the SON. It is plausible that HSD also induces changes in astrocytes, potentially triggering the expression of “eat me” tags on the astrocyte surface, marking them for elimination by microglia, or reducing the expression of “don’t eat me” signals that would normally protect them from phagocytosis^61,62^.

A recent study found that increased endocannabinoid tone during development promotes microglia phagocytosis of astrocytic precursors, influencing newborn astrocyte numbers and neurocircuit maturation in the amygdala^63^. Since magnocellular neurons release endocannabinoids in an activity-dependent manner^64^, HSD-induced increases in magnocellular VP neuron activity may trigger local endocannabinoid release, promoting microglial phagocytosis of astrocytic processes. Moreover, as endocannabinoids can act as chemoattractants, inducing microglia migration^65^, they may mediate microglia accumulation, activation, and engulfment of astrocytic processes around magnocellular VP neurons in HSD.

Many other molecules are upregulated or downregulated in response to HSD, and various candidate signaling molecules or their combinations may be involved in this process. Further experiments are needed to decipher the molecular mechanisms underlying microglial activation in HSD.

### Microglia-mediate pruning of astrocyte processes in high-salt diet

Astrocytes play a central role in sculpting neuronal networks, modulating synaptic transmission, neurotransmitter clearance, and synaptic plasticity^3^. Thus, structural remodeling of the astrocyte network in HSD can modulate neural activity in multiple ways^4^. Previous ultrastructural studies characterised this remarkable structural plasticity of astrocytes and proposed that decreased astrocytic coverage in HSD is mediated by astrocyte “withdrawal” or “retraction”^5,7,8,16,18,19,22^. These studies utilized ultrastructural examination with electron microscopy, allowing for the evaluation and quantification of specific details related to neuronal appositions and synaptic number. However, they lack evidence supporting the notion that astrocytes “retract” their processes. We replicated these studies by examining this phenomenon using immunofluorescence, that allows for a more comprehensive analysis of the spatial organization of different cell populations and the quantification of multiple protein components.

Our data reveal that the decrease in astrocyte coverage is associated with a reduction in multiple astrocyte proteins, including GFAP, S100β, ALDH1L1, GLT1, and vimentin (Fig. 1). These changes are associated with a gradual decrease in astrocytic material in the SON and PVN, which can be quantified and corroborated by immunofluorescence and by analyzing reductions in total SON astrocytic protein levels in HSD-fed rats (Fig. S1). Importantly, this reduction is not a result of astrocyte death but rather a decrease in the density of astrocyte processes, since the total number of SON and PVN astrocytes is not affected by the HSD (Fig. S2C and S2D). Notably, we demonstrated that these changes are mediated by reactive microglia that specifically phagocytose astrocyte material (Fig. 4 and Videos S1 & S2), leading to the reduction in astrocyte coverage. Thus, our data suggest that the decrease in astrocytic coverage in HSD represents microglia-mediated pruning of astrocytes rather than astrocyte retraction.

### Microglia-mediated reduction of astrocytic processes underlies changes in VP neuron activity

While significant changes in astrocyte coverage can affect neuronal excitation through various mechanisms^40,41,66^, we examined the hypothesis that decreased astrocytic coverage causes an impaired synaptic glutamate clearance, leading to glutamate spillover into the extracellular space, thereby activating extrasynaptic glutamate receptors and increasing tonic activation of VP neurons. Our results demonstrate that while in control condition, blocking extrasynaptic NMDARs does not affect the spontaneous firing activity of SON VP neurons, inhibiting these receptors in the SON from HSD-fed rats decreases the firing rate of VP neurons, revealing the presence of a tonic excitatory current in this condition. Furthermore, inhibiting microglia-mediated pruning of astrocytes in HSD restores this effect, suggesting that the dense network of astrocytic processes around VP neurons is essential to prevent the tonic activation of extrasynaptic NMDARs. Our findings suggest that the effectiveness of astrocytes in controlling VP neuron activity and preventing their activation by extrasynaptic NMDARs is mediated by the major glutamate transporter GLT1 (also known as excitatory amino acid transporter 2, or EAAT2). Blocking this transporter increases the spontaneous firing of SON VP neurons in control conditions, mimicking the stimulating effect of HSD on their activity. This also suggests that extrasynaptic NMDARs are normally present on the surface of VP neurons under control conditions but remain inactive due to the efficient removal of extrasynaptic glutamate by astrocytes. Our findings imply that the regulation of VP neuron firing by astrocytes (via the astrocytic transporter GLT1) is controlled by microglia. Blocking microglia activation and proliferation by inhibiting CSF1R-mediated signaling prevents microglia-mediated pruning of astrocytic processes (Figs. 3 & S5-7), diminishes the reduction in GLT1 levels (Fig. S9C) and GLT1 function (Fig. 5C), and confines the spontaneous firing of VP neurons to control levels (Fig. 6J).

Finally, we showed that interactions between neurons, microglia, and astrocytes, determine neuronal activity levels under normal conditions and underlie functional state-dependent plasticity, facilitating VP neuron firing and systemic VP release, which contribute to salt-dependent hypertension. Inhibiting microglia activation not only prevents structural plasticity of astrocytes and increased excitation of VP neurons, but also significantly reduces HSD-induced increases in blood pressure and attenuates salt-dependent hypertension (Fig. 7C & 7D).

Since previous studies have demonstrated that increased VP secretion contributes to elevated blood pressure in various models of salt-dependent hypertension^38,67,68^, we primarily focused on magnocellular VP neurons. However, given that oxytocin is a natriuretic hormone in rats, nearby oxytocin neurons may also be affected by HSD, providing an adaptive mechanism to facilitate sodium excretion and reduce the impact of HSD. Therefore, it is important to investigate whether microglia-mediated pruning of astrocytes also drives changes in oxytocin neuron activity in HSD, or other conditions associated with structural astrocyte plasticity in the magnocellular hypothalamo-neurohypophysial system.

In summary, our study reveals the capacity of microglia to phagocytose and prune astrocytic processes in the adult brain. These findings uncover a novel mechanism by which microglia can regulate synaptic transmission and neuronal activity via structural remodeling of the astrocytic network. Microglia density and morphology vary between brain regions^69,70^, hence it remains unclear whether the capacity of microglia to remodel astrocytes is unique to the specific hypothalamic region. Future studies should examine whether a similar mechanism can underlie additional forms of structural plasticity in other brain regions under certain physiological or pathological conditions.

## Acknowledgments

This work was supported by the Canadian Institutes of Health Research Project Grants PJT-153009 & 178195 (MPK). MPK is a recipient of a Heart & Stroke Foundation of Canada National New Investigator Award.

## Author contributions

MPK designed the research program and experiments. NG, OM, CL, CQC, BL, GL, JY, CW, and MPK conducted the experiments. NG, OM, CL, CQC, PMC, BL, JY, CW, JF, BS, SPS, MH, and MPK analyzed the data. MPK wrote the manuscript, and NG, KYC, AK, CWB, and MPK reviewed and edited the manuscript.

---

**Fig. S1:**
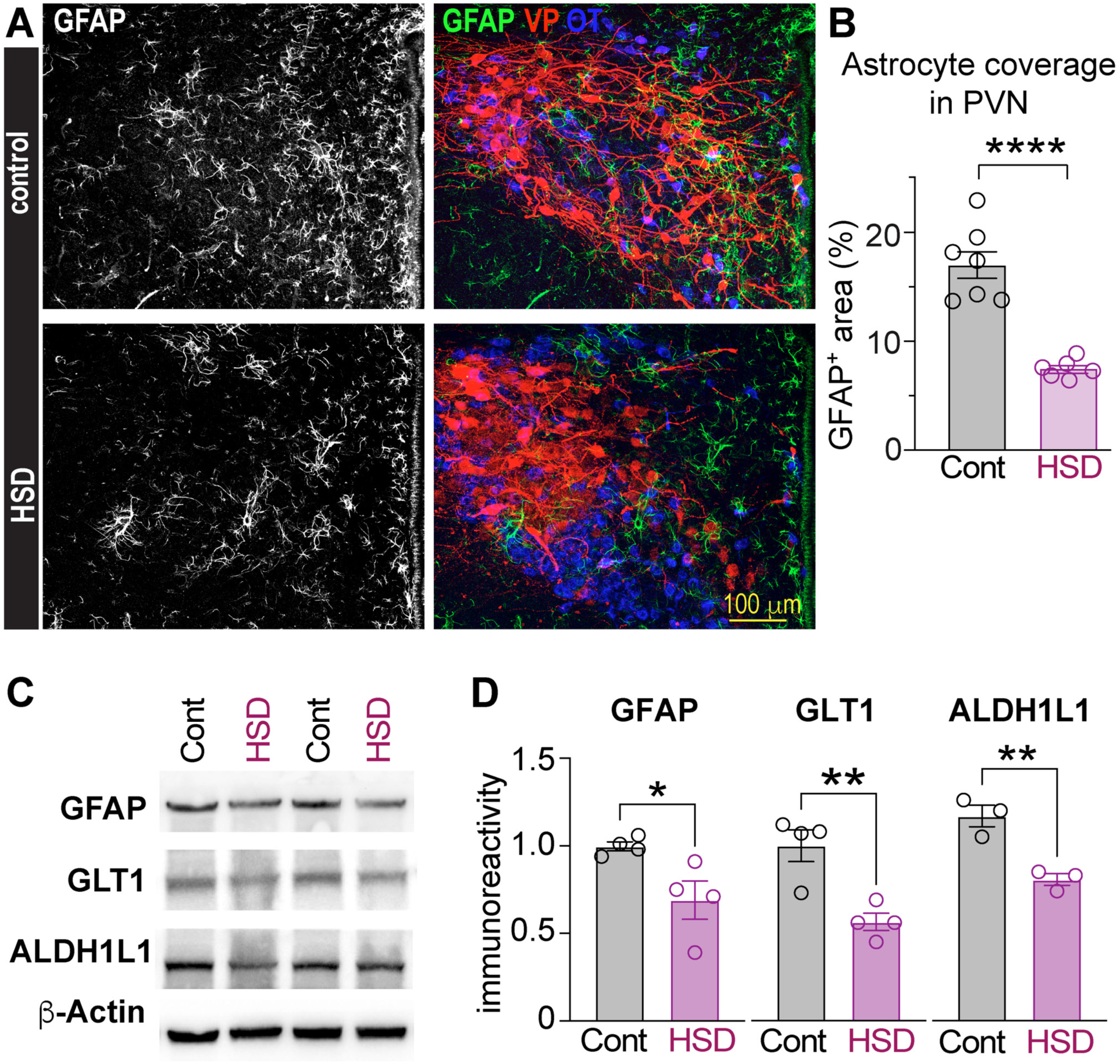
Decreased astrocyte coverage in high-salt diet is illustrated by reduced levels of astrocytic markers. (**A**) Coronal rat brain sections immunolabeled to visualize astrocytes (GFAP, white or green) and neurons secreting VP (red) or oxytocin (blue) in the PVN of control rats (top) or rats fed HSD (bottom). (**B**) Astrocyte coverage is analyzed as GFAP-immunopositive PVN area in rats fed control or HSD (n=7 control, n=6 HSD rats, Student’s t-test). (**C**) Western blot on homogenized SON of rats fed control or HSD. Membrane was cut and probed for astrocytic markers: GFAP, GLT1, and ALDH1L1, with β-actin as a loading control. All images are from the same membrane. (**E**) Quantifications of GFAP, GLT1, and ALDH1L1 bands from (**D**), normalized to the respective β-actin band in that lane (n=3-4 rats/group, Student’s t-test). Each data point represents one rat, \*\*\*\**P* < 0.0001, \*\**P* < 0.001, \*\**P* < 0.01, \**P* < 0.05, ns, not significant.

**Fig. S2:**
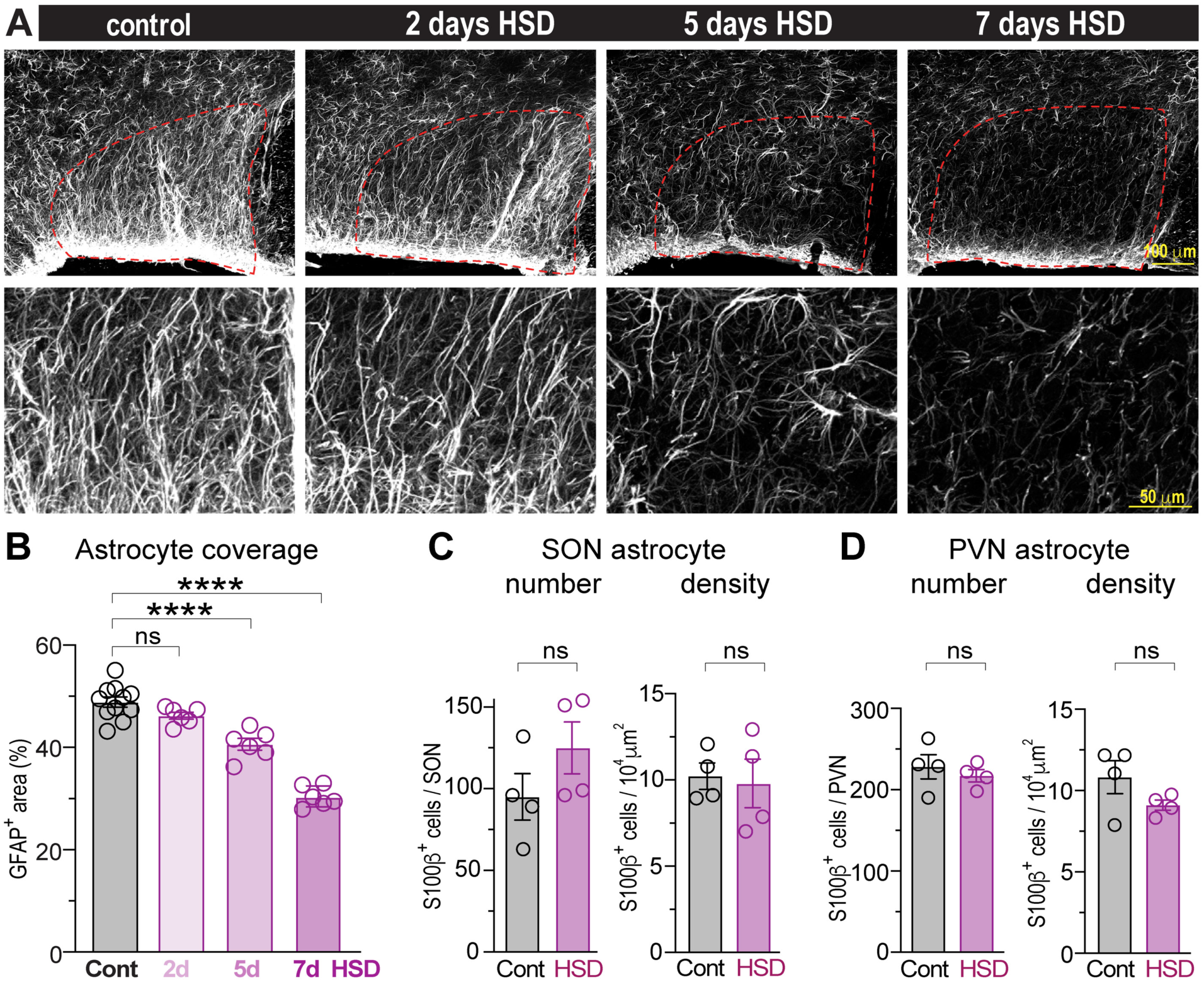
High-salt diet induces a gradual reduction in astrocytic coverage. (**A**) Coronal rat brain sections immunolabeled for GFAP to visualize astrocytes in the SON of control rats or rats fed HSD for 2, 5, or 7 days. Red dotted lines outline VP neuron location on top images. Bottom images show magnified central VP-containing SON areas corresponding to the top image. (**B**) Astrocyte coverage is analyzed as GFAP-immunopositive SON area in rats fed control diet or following 2, 5, or 7 days of HSD (n=10 control rats, n=6 rats/group for 2, 5, and 7 days of HSD, one-way ANOVA with Tukey’s post-hoc test). (**C**, **D**) Quantification of the total number (left) and density (right) of S100β-positive astrocytes in the SON (**C**) and PVN (**D**) of rats fed control diet or 7 days of HSD (n=4 rats/group, student t-test). Each data point represents one rat, \*\*\*\**P* < 0.0001, ns, not significant.

**Fig. S3:**
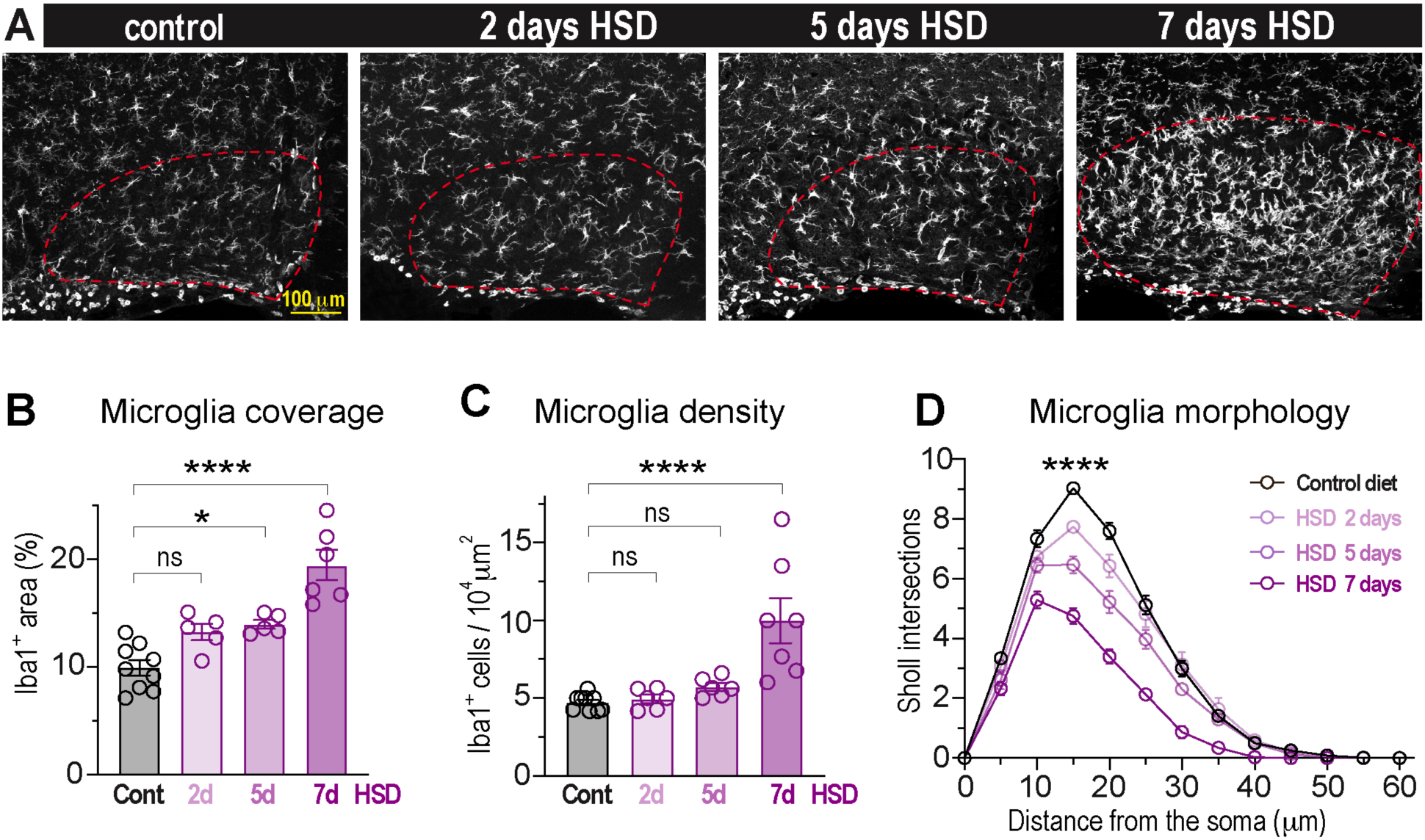
High-salt diet induces gradual changes in microglia density and morphology. (**A**) Coronal rat brain sections immunolabeled for Iba1 to visualize microglia in the SON of control rats or rats fed HSD for 2, 5, or 7 days. Red dotted lines outline VP neuron location. Quantification of microglia coverage, analyzed as Iba1-immunopositive SON area (**B**), and microglia density (**C**) in rats fed control diet for 2, 5, or 7 days of HSD (n=9 control rats, n=5 rats/group for 2 and 5 days, and n=6 for 7 days of HSD, one-way ANOVA with Tukey’s post-hoc test). (**D**) Distribution of Sholl intersection number by the distance from microglia soma in the SONs in control rats and rats fed HSD for 2, 5, or 7 days, (n=5 rats/group, 20 cells per animal, p < 0.0001 by two-way ANOVA with Tukey’s post-hoc test). Data presented as mean ± SEM, \**P* < 0.05, \*\*\*\**P* < 0.0001, ns, not significant; each data point represents one rat.

**Fig. S4:**
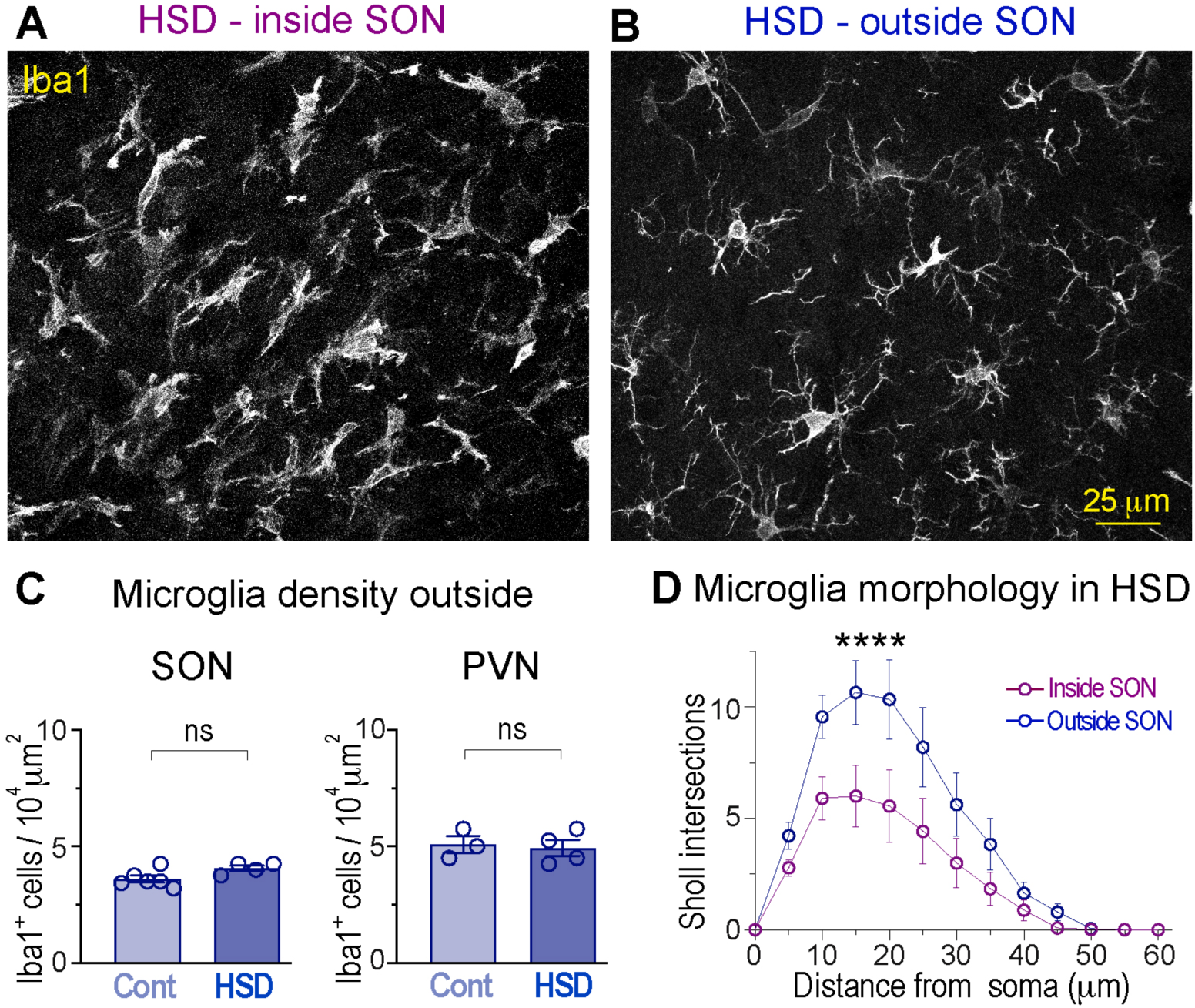
High-salt diet-induced activation of microglia inside but not outside of the SON. Coronal rat brain sections from rats fed 7 days HSD immunolabeled to visualize microglia (Iba1) inside the SON (**A**) and in the adjacent area, immediately outside the SON (**B**). (**C**) Microglia density in the areas adjacent to the SON and PVN in rats fed control diet or HSD (n=6 control and n=4 HSD rats for SON; n=3 control and n=4 HSD rats for PVN, Student’s t-test, ns, not significant). (**D**) Distribution of Sholl intersection number by the distance from microglia soma found inside and outside the SON from HSD-fed rats (n=5 rats, 20 cells/area/animal), \*\*\*\**P* < 0.0001, by two-way ANOVA with Tukey’s post-hoc test. Data presented as mean ± SEM, each data point represents one rat.

**Fig. S5:**
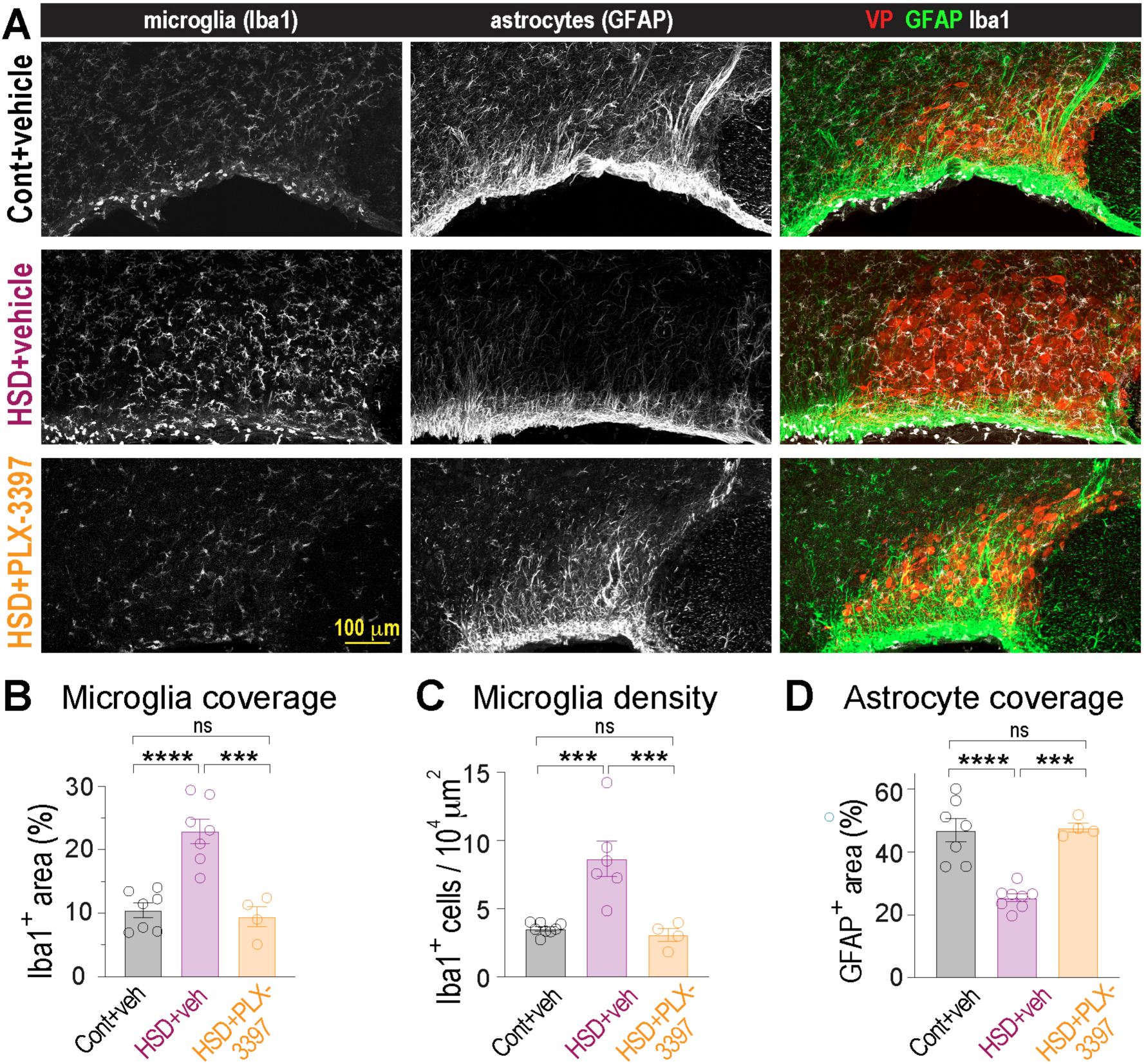
Microglia inhibition with PLX3397 blocks astrocyte remodeling in high-salt diet. (**A**) SON sections from control (cont) and HSD-fed rats treated with i.p. vehicle (veh), and HSD rats treated with 30 mg/kg/day PLX3397 i.p. labeled with markers of microglia (Iba1), astrocytes (GFAP), and VP neurons. Quantification of microglia coverage, analyzed as Iba1-immunopositive SON area (**B**), microglia density (**C**), and astrocyte coverage, analyzed as GFAP-immunopositive SON area (**D**), in rats exposed to different treatments (n=7 rats/group for cont+veh and HSD+veh, and n=4 rats for HSD + PLX3397, one-way ANOVA with Tukey’s post-hoc test). Data presented as mean ± SEM, each data point represents one rat, \*\*\*\**P* < 0.0001, \*\*\**P* < 0.001, ns, not significant.

**Fig. S6:**
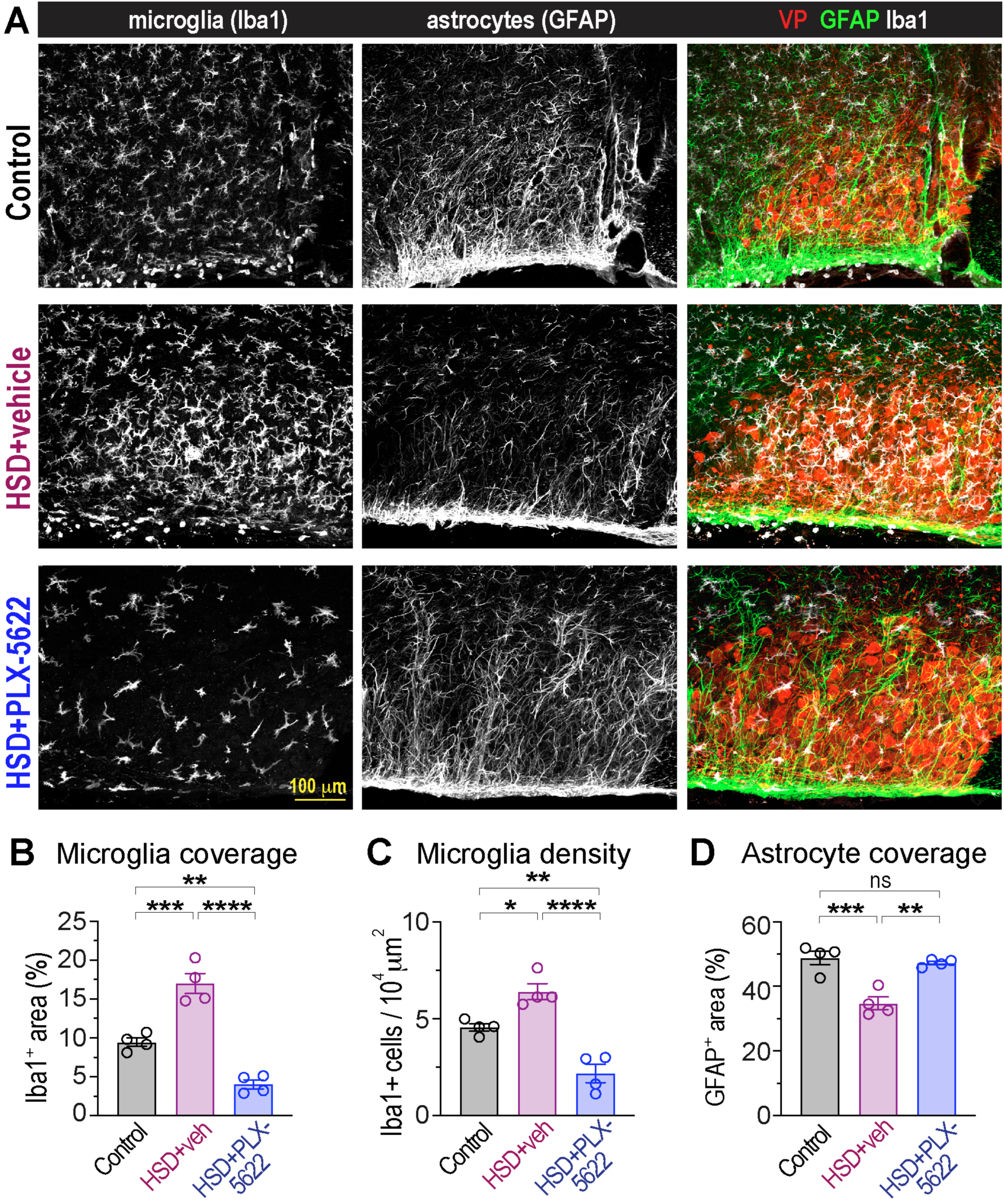
Microglia inhibition with PLX5622 blocks astrocyte remodeling in high-salt diet. (**A**) SON sections from control and HSD-fed rats treated with i.p. vehicle, and HSD rats treated with 50 mg/kg PLX5622 i.p. twice per day labeled with markers of microglia (Iba1), astrocytes (GFAP), and VP neurons. Quantification of microglia coverage, analyzed as Iba1-immunopositive SON area (**B**), microglia density (**C**), and astrocyte coverage, analyzed as GFAP-immunopositive SON area (**D**), in rats exposed to different treatments (n = 4 rats/group, one-way ANOVA with Tukey’s post-hoc test). Data presented as mean ± SEM, each data point represents one rat, \*\*\*\**P* < 0.0001, \*\*\**P* < 0.001, \*\**P* < 0.01, \**P* < 0.05, ns, not significant.

**Fig. S7:**
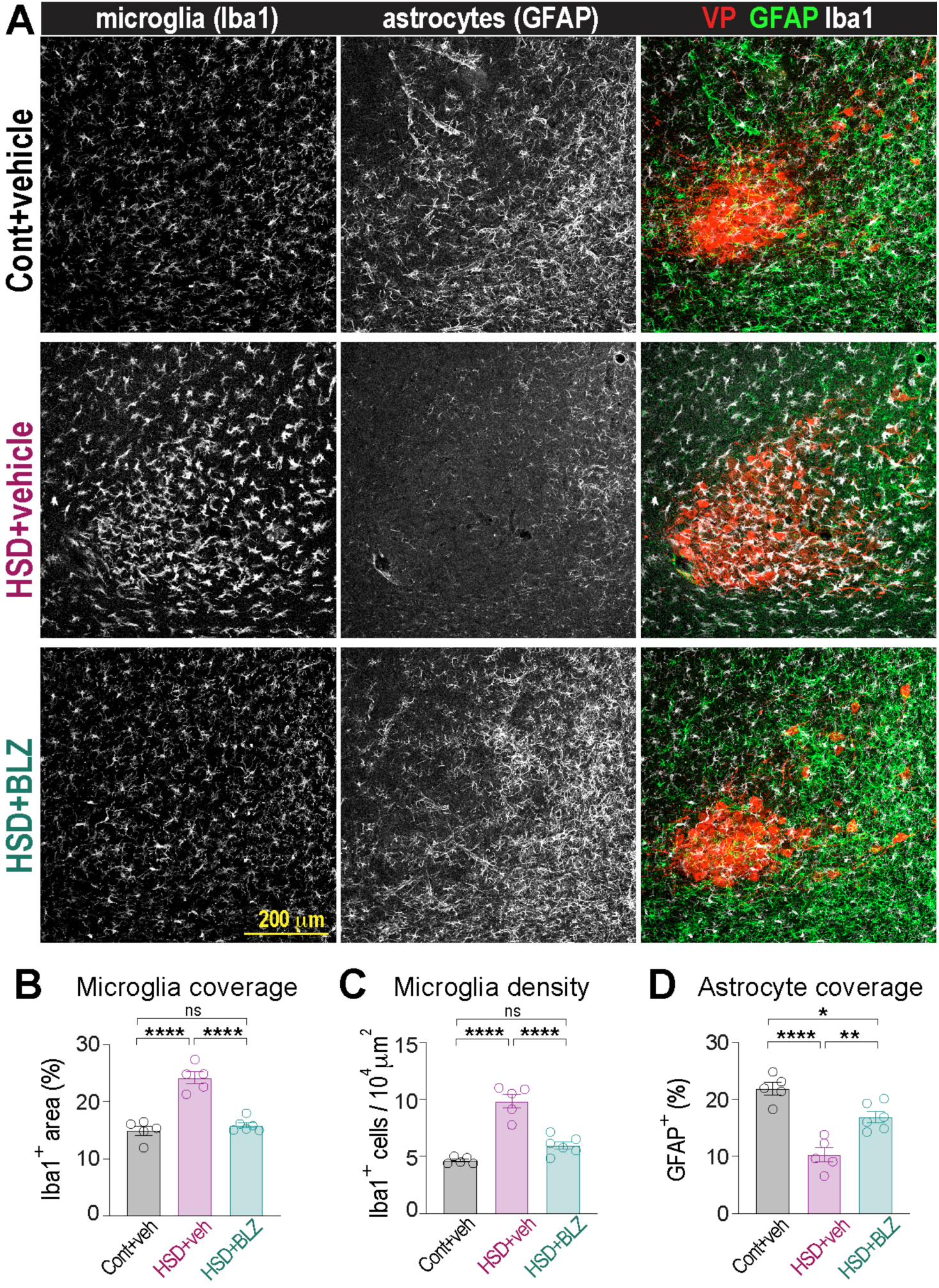
Microglia inhibition with BLZ945 blocks astrocyte remodeling in PVN from high-salt diet. (**A**) PVN sections from control (cont) and HSD-fed rats treated i.c.v. with vehicle (veh) or 0.3 mg/kg/day BLZ945, labeled with markers of microglia (Iba1), astrocytes (GFAP), and VP neurons. Quantification of microglia coverage, analyzed as Iba1-immunopositive PVN area (**B**), microglia density (**C**), and astrocyte coverage, analyzed as GFAP-immunopositive PVN area (**D**), in rats exposed to different treatments (n=5 rats/group for cont+veh and HSD+veh, and n=6 rats for HSD + BLZ, one-way ANOVA with Tukey’s post-hoc test). Data presented as mean ± SEM, each data point represents one rat, \*\*\*\**P* < 0.0001, \*\*\**P* < 0.001, \*\**P* < 0.01, \**P* < 0.05, ns, not significant.

**Fig. S8:**
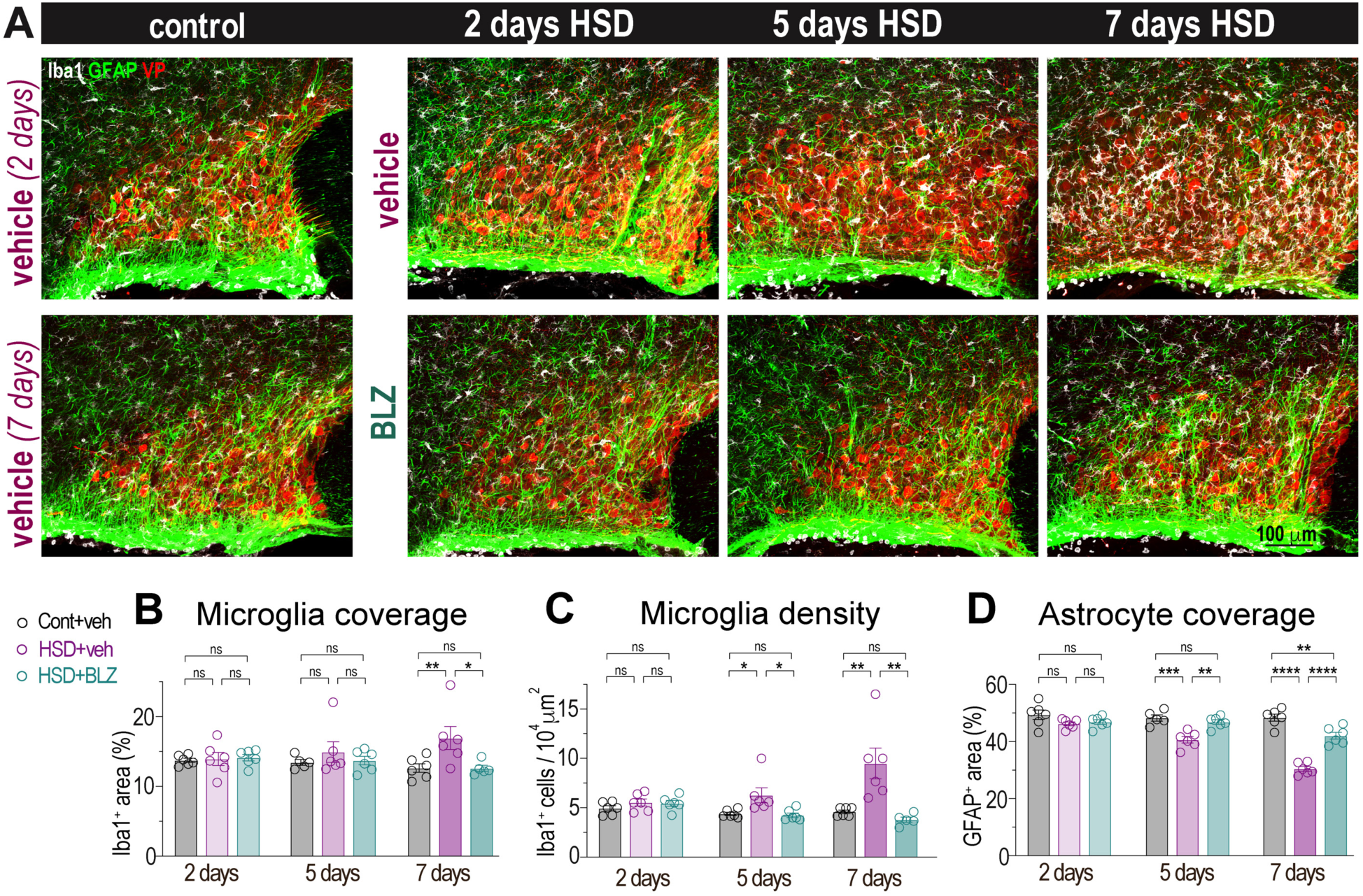
Microglia inhibition with BLZ945 prevents accumulation of microglia and astrocyte remodeling in high-salt diet. Rats treated with vehicle or 0.3 mg/kg/day BLZ945 i.c.v. were subjected to control diet or HSD for 2, 5, or 7 days. (**A**) SON sections from different groups are labeled with markers of microglia (Iba1, white), astrocytes (GFAP, green), and VP neurons (VP, red). Quantification of microglia coverage, analyzed as Iba1-immunopositive SON area (**B**), microglia density (**C**), and astrocyte coverage, analyzed as GFAP-immunopositive SON area (**D**), in rats exposed to different treatments (n=5-6 rats/group, one-way ANOVA with Tukey’s post-hoc test). Data presented as mean ± SEM, each data point represents one rat, \*\*\*\**P* < 0.0001, \*\*\**P* < 0.001, \*\**P* < 0.01, \**P* < 0.05, ns, not significant.

**Fig. S9:**
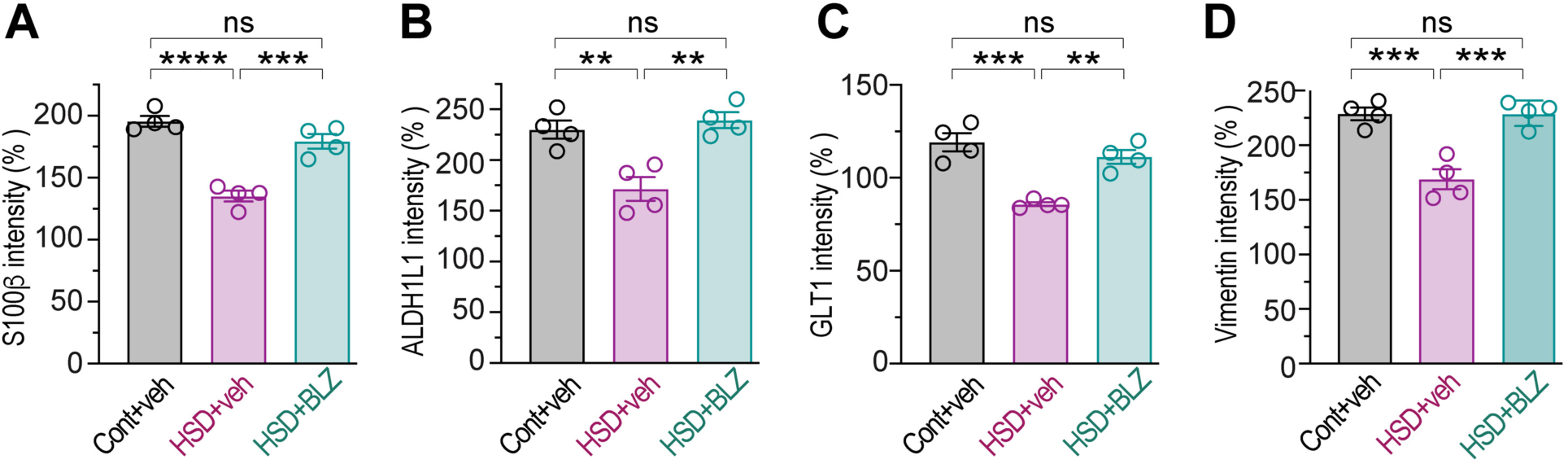
Microglia inhibition with BLZ945 blocks high-salt diet-induced decreases in astrocyte markers in the SON. Rats were fed control diet or HSD and treated i.c.v. with vehicle (veh) or 0.3 mg/kg/day BLZ945, labeled with astrocyte markers (S100β, ALDH1L1, GLT1, and vimentin) and VP neurons. Quantifications of the relative levels of S100β (**α**), ALDH1L1 (**B**), GLT1 (**C**), and vimentin (**D**) in VP-positive SON area in rats exposed to different treatments (n=4 rats/group, one-way ANOVA with Tukey’s post-hoc test). Data presented as mean ± SEM, each data point represents one rat, \*\*\*\**P* < 0.0001, \*\*\**P* < 0.001, \*\**P* < 0.01, ns, not significant.

**Fig. S10:**
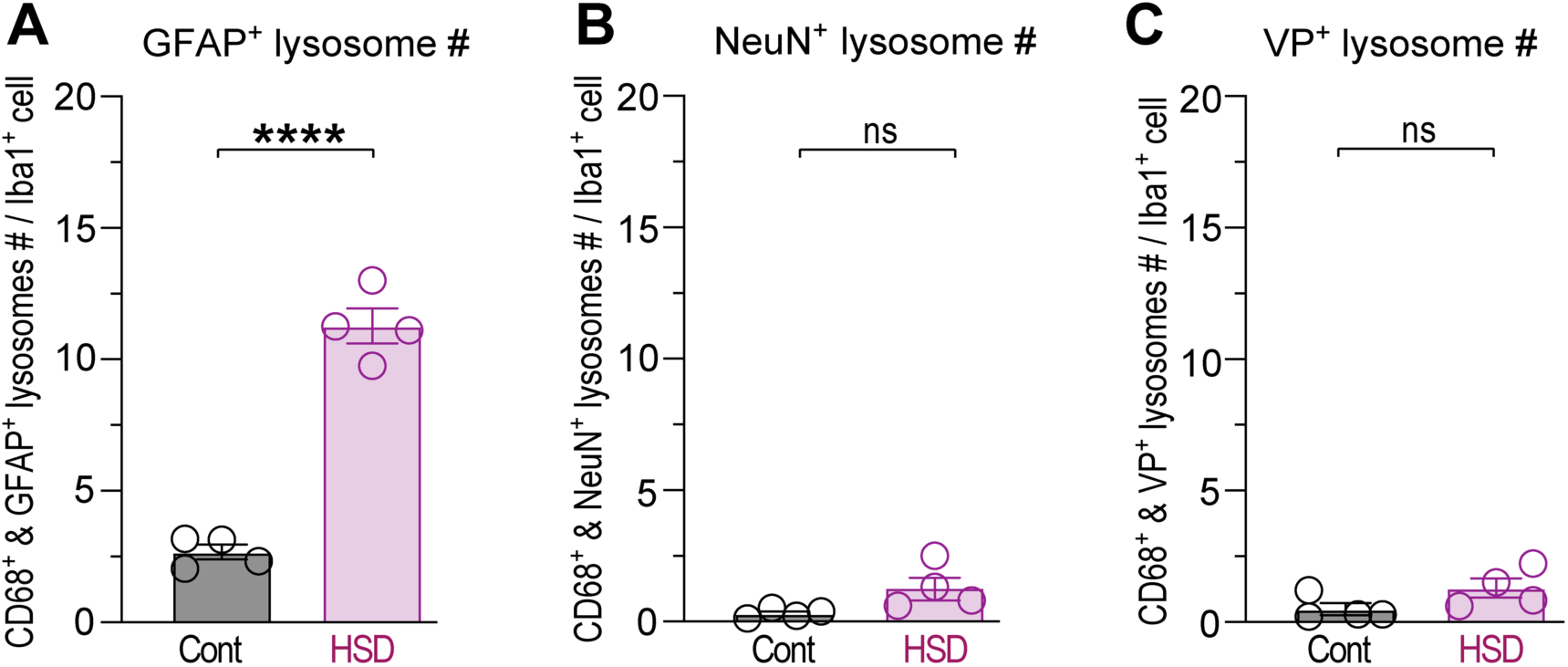
Microglia increase phagocytosis of astrocytic but not neuronal elements in high-salt diet. SON sections from control and HSD-fed rats were stained for Iba1 to identify microglia, microglia-specific lysosomal marker (CD68), astrocytic marker GFAP, and either general pan-neuronal marker (NeuN) or a marker of VP neurons (VP) and analyzed with Super-resolution Airyscan. Bar graphs show the mean number of CD68+ lysosomes containing GFAP (**A**), NeuN (**B**), or VP (**C**) in control and HSD-treaded rats. Each data point represents one rat, 15-20 microglia were analyzed per rat, n=4 rats/group, compared by Student’s t-test. All data presented as mean ± SEM, \*\*\*\**P* < 0.0001, ns, not significant.

**Fig. S11:**
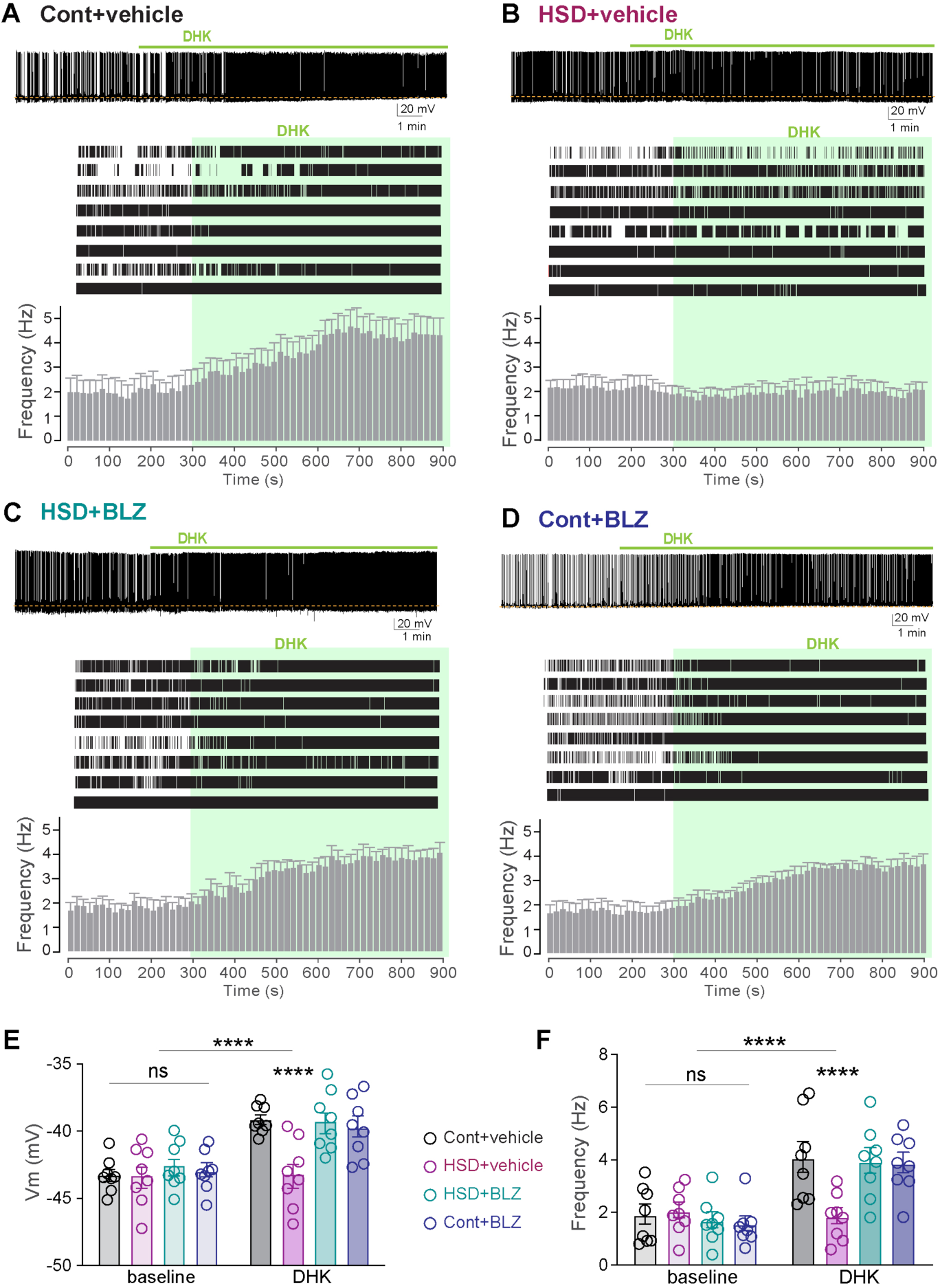
Microglia-mediated astrocyte pruning impairs synaptic glutamate clearance by glutamate transporter. Top: Sample current-clamp traces show the effect of bath application of glial glutamate transporter blocker DHK (100 µM, green bar) on action potential firing of VP neurons in SON brain slices prepared from vehicle-treated control (**A**) and HSD (**B**) rats, and BLZ945-treated HSD (**C**) and control (**D**) rats. Dashed lines show membrane potential at baseline. Middle: Current-clamp recordings showing changes in action potential firing in 8 rats in response to DHK application (green shaded area). Bottom: Bar graphs show mean ± SEM firing frequency changes corresponding to above 8 traces before and after the application of DHK (green). Bar graphs show membrane potential (**E**) and firing rate (**F**) at baseline and after DHK application in all groups (n=8 rats/group, repeated measures two-way ANOVA with Tukey’s multiple comparisons post-hoc test). Data presented as mean ± SEM, each data point represents one rat, **** *P* < 0.0001, ns, not significant.

**Fig. S12:**
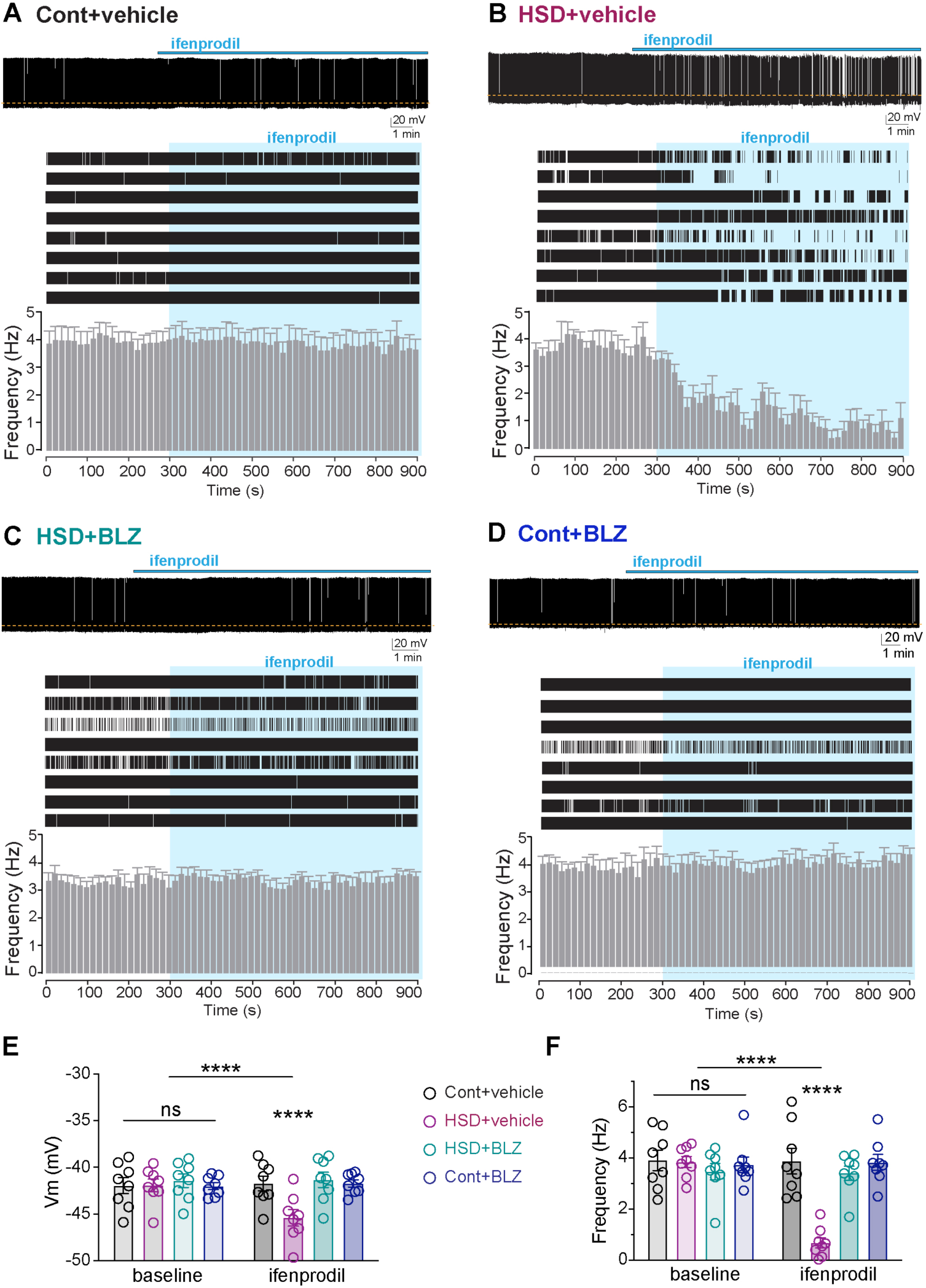
Microglia-mediated astrocyte pruning reveals the presence of tonic excitatory extrasynaptic NMDAR currents in VP neurons. Top: Sample current-clamp traces show the effect of bath application of the selective blocker of extrasynaptic NR2B NMDAR subunit, ifenprodil (10 µM, blue bar) on action potential firing of VP neurons in SON brain slices prepared from vehicle-treated control (**A**) and HSD (**B**) rats, and BLZ945-treated HSD (**C**) and control (**D**) rats. Dashed lines show membrane potential at baseline. Middle: Current-clamp recordings showing changes in action potential firing in 8 rats in response to ifenprodil application (blue shaded area). Bottom: Bar graphs show mean ± SEM firing frequency changes corresponding to above 8 traces before and after the application of ifenprodil (blue). Bar graphs show membrane potential (**E**) and firing rate (**F**) at baseline and after ifenprodil application in all groups (n=8 rats/group, repeated measures two-way ANOVA with Tukey’s multiple comparisons post-hoc test). Data presented as mean ± SEM, each data point represents one rat, \*\*\*\**P* < 0.0001, ns, not significant.

## Supplemental Videos

**Video <u>S1</u>:** IMARIS 3D reconstruction of a microglia from the SON of rat fed control diet and injected with vehicle. Microglial surface reconstruction from the Iba1 channel is shown in purple. Reconstructed surfaces of lysosomes from the CD68 channel are shown in white.

**Video <u>S2</u>:** IMARIS 3D reconstruction of a microglia from a rat fed HSD and treated with the vehicle. Microglial surface reconstruction from the Iba1 channel is shown in purple. Reconstructed surfaces of lysosomes from the CD68 channel are shown in white. Reconstructed surfaces of GFAP and S100β inside the lysosomes are shown in orange and green respectively.

**Video <u>S3</u>:** IMARIS 3D reconstruction of a microglia from a rat fed HSD and injected with BLZ. Reconstructed surfaces of lysosomes from the CD68 channel are shown in white.

## Supplemental Tables

**Table S1:** Drug concentrations, targets, and routes of administration

| short name | alternative name | concentration | main target | references |
| --- | --- | --- | --- | --- |
| PLX3397 | Pexidartinib | 30 mg/kg/day i.p. | Colony stimulating factor 1 receptor (CSF1R) | 29, 71 |
| PLX5622 | PLX5622 | 100 mg/kg/day i.p. | Colony stimulating factor 1 receptor (CSF1R) | 28, 72 |
| BLZ945 | Sotuletinib | 0.3 mg/kg/day i.c.v. | Colony stimulating factor 1 receptor (CSF1R) | 73, 33 |
| DHK | Dihydrokainate | 100 mM | Glutamate transporter 1 (GLT1) | 74 |
| Ifenprodil | Ifenprodil | 10 μM | NR2B NMDA receptors | 75 |
| AP-5 | (2R)-amino-5-phosphonovaleric acid | 100 μM | NMDA receptors | 76 |
| KYN | Kynurenic acid | 1 mM | Ionotropic glutamate receptors (AMPA, NMDA, and kainite) | 77 |
| TTX | Tetrodotoxin | 1 nM | Voltage-gated Na<sup>+</sup> channels | 78 |

**Table S2:** Statistical analyses

| Figure & panel | Compared groups | Statistical test | Post-hoc test (for multiple comparison) | P value (adjusted) | P value (summary) |
| --- | --- | --- | --- | --- | --- |
| Fig. 1F | Cont vs HSD | unpaired 2-tailed t-test |  | <0.0001 | **** |
| Fig. 1G | Cont vs HSD | unpaired 2-tailed t-test |  | 0.000065 | **** |
| Fig. 1H | Cont vs HSD | unpaired 2-tailed t-test |  | 0.007454 | ** |
| Fig. 1I | Cont vs HSD | unpaired 2-tailed t-test |  | 0.000562 | *** |
| Fig. 1J | Cont vs HSD | unpaired 2-tailed t-test |  | 0.001490 | ** |
| Fig. 2E | Cont vs HSD | unpaired 2-tailed t-test |  | <0.0001 | **** |
| Fig. 2F | Cont vs HSD | unpaired 2-tailed t-test |  | 0.000698 | *** |
| Fig. 2G | Cont vs HSD | unpaired 2-tailed t-test |  | 0.000161 | *** |
| Fig. 2H | Cont vs HSD | 2-way ANOVA | Šídák's | <0.0001 | **** |
| Fig. 3B |  | 1-way ANOVA | Tukey's | <0.0001 | **** |
|  | Cont+veh vs HSD+veh |  |  | <0.0001 | **** |
|  | Cont+veh vs HSD+BLZ |  |  | 0.5731 | ns |
|  | HSD+veh vs HSD+BLZ |  |  | <0.0001 | **** |
| Fig. 3C |  | 1-way ANOVA | Tukey's | <0.0001 | **** |
|  | Cont+veh vs HSD+veh |  |  | <0.0001 | **** |
|  | Cont+veh vs HSD+BLZ |  |  | 0.9441 | ns |
|  | HSD+veh vs HSD+BLZ |  |  | <0.0001 | **** |
| Fig. 3D |  | 2-way ANOVA | Tukey's | <0.0001 | **** |
|  | Cont+veh vs HSD+veh |  |  | <0.0001 | **** |
|  | Cont+veh vs HSD+BLZ |  |  | 0.0001 | *** |
|  | HSD+veh vs HSD+BLZ |  |  | <0.0001 | **** |
| Fig. 3E |  | 1-way ANOVA | Tukey's | <0.0001 | **** |
|  | Cont+veh vs HSD+veh |  |  | <0.0001 | **** |
|  | Cont+veh vs HSD+BLZ |  |  | 0.0002 | *** |
|  | HSD+veh vs HSD+BLZ |  |  | <0.0001 | **** |
| Fig. 4C |  | 1-way ANOVA | Tukey's | <0.0001 | **** |
|  | Cont+veh vs HSD+veh |  |  | <0.0001 | **** |
|  | Cont+veh vs HSD+BLZ |  |  | 0.0160 | * |
|  | HSD+veh vs HSD+BLZ |  |  | <0.0001 | **** |
| Fig. 4D |  | 1-way ANOVA | Tukey's | <0.0001 | **** |
|  | Cont+veh vs HSD+veh |  |  | <0.0001 | **** |
|  | Cont+veh vs HSD+BLZ |  |  | 0.1128 | ns |
|  | HSD+veh vs HSD+BLZ |  |  | <0.0001 | **** |
| Fig. 4E |  | 1-way ANOVA | Tukey's | <0.0001 | **** |
|  | Cont+veh vs HSD+veh |  |  | <0.0001 | **** |
|  | Cont+veh vs HSD+BLZ |  |  | 0.0370 | * |
|  | HSD+veh vs HSD+BLZ |  |  | <0.0001 | **** |
| Fig. 4F |  | 1-way ANOVA | Tukey's | <0.0001 | **** |
|  | Cont+veh vs HSD+veh |  |  | <0.0001 | **** |
|  | Cont+veh vs HSD+BLZ |  |  | 0.0274 | * |
|  | HSD+veh vs HSD+BLZ |  |  | <0.0001 | **** |
| Fig. 5E |  | 2-way RM ANOVA | Šídák's | <0.0001 | **** |
|  | Cont+veh: baseline vs DHK |  |  | <0.0001 | **** |
|  | HSD+veh: baseline vs DHK |  |  | 0.9980 | ns |
|  | HSD+BLZ: baseline vs DHK |  |  | <0.0001 | **** |
|  | Cont+BLZ: baseline vs DHK |  |  | <0.0001 | **** |
| Fig. 5F |  | 2-way RM ANOVA | Šídák's | <0.0001 | **** |
|  | Cont+veh, baseline vs DHK |  |  | <0.0001 | **** |
|  | HSD+veh: baseline vs DHK |  |  | 0.7998 | ns |
|  | HSD+BLZ: baseline vs DHK |  |  | <0.0001 | **** |
|  | Cont+BLZ: baseline vs DHK |  |  | <0.0001 | **** |
| Fig. 6E |  | 1-way RM ANOVA | Tukey's | 0.0005 | *** |
|  | Cont+veh vs HSD+veh |  |  | 0.0009 | *** |
|  | Cont+veh vs HSD+BLZ |  |  | 0.9686 | ns |
|  | Cont+veh vs Cont+BLZ |  |  | 0.8790 | ns |
|  | HSD+veh vs HSD+BLZ |  |  | 0.0021 | ** |
|  | HSD+veh vs Cont+BLZ |  |  | 0.0039 | ** |
|  | HSD+BLZ vs Cont+BLZ |  |  | 0.9911 | ns |
| Fig. 6F |  | 1-way RM ANOVA | Tukey's | 0.3999 | ns |
|  | Cont+veh vs HSD+veh |  |  | 0.8574 | ns |
|  | Cont+veh vs HSD+BLZ |  |  | 0.6558 | ns |
|  | Cont+veh vs Cont+BLZ |  |  | 0.8049 | ns |
|  | HSD+veh vs HSD+BLZ |  |  | 0.6011 | ns |
|  | HSD+veh vs Cont+BLZ |  |  | 0.7953 | ns |
|  | HSD+BLZ vs Cont+BLZ |  |  | 0.8405 | ns |
| Fig. 6G |  | 1-way RM ANOVA | Tukey's | 0.2086 | ns |
|  | Cont+veh vs HSD+veh |  |  | 0.8914 | ns |
|  | Cont+veh vs HSD+BLZ |  |  | 0.6046 | ns |
|  | Cont+veh vs Cont+BLZ |  |  | 0.7247 | ns |
|  | HSD+veh vs HSD+BLZ |  |  | 0.2426 | ns |
|  | HSD+veh vs Cont+BLZ |  |  | 0.3280 | ns |
|  | HSD+BLZ vs Cont+BLZ |  |  | 0.9969 | ns |
| Fig. 6L |  | 2-way RM ANOVA | Šídák's | <0.0001 | **** |
|  | Cont+veh: baseline vs ifenpr |  |  | 0.9558 | ns |
|  | HSD+veh: baseline vs ifenpr |  |  | <0.0001 | **** |
|  | HSD+BLZ: baseline vs ifenpr |  |  | 0.9978 | ns |
|  | Cont+BLZ: baseline vs ifenpr |  |  | 0.9132 | ns |
| Fig. 6M |  | 2-way RM ANOVA | Šídák's | <0.0001 | **** |
|  | Cont+veh, baseline vs ifenpr |  |  | <0.0001 | **** |
|  | HSD+veh: baseline vs ifenpr |  |  | 0.7998 | ns |
|  | HSD+BLZ: baseline vs ifenpr |  |  | <0.0001 | **** |
|  | Cont+BLZ: baseline vs ifenpr |  |  | <0.0001 | **** |
| Fig. 7C |  | 1-way ANOVA | Tukey's | <0.0001 | **** |
|  | Cont+veh vs HSD+veh |  |  | <0.0001 | **** |
|  | Cont+veh vs HSD+PLX |  |  | 0.0018 | ** |
|  | HSD+veh vs HSD+PLX |  |  | 0.0176 | * |
| Fig. 7D |  | 1-way ANOVA | Tukey's | <0.0001 | **** |
|  | Cont+veh vs HSD+veh |  |  | <0.0001 | **** |
|  | Cont+veh vs HSD+BLZ |  |  | 0.2034 | ns |
|  | HSD+veh vs HSD+BLZ |  |  | <0.0001 | **** |
| Fig. S1B | Cont vs HSD | unpaired 2-tailed t-test |  | <0.0001 | **** |
| Fig. S1D | GFAP: Cont vs HSD | unpaired 2-tailed t-test |  | 0.0333 | * |
| Fig. S1D | GLT1: Cont vs HSD | unpaired 2-tailed t-test |  | 0.0056 | ** |
| Fig. S1D | ADL1H1: Cont vs HSD | unpaired 2-tailed t-test |  | 0.0069 | ** |
| Fig. S2B |  | 1-way ANOVA | Tukey's | <0.0001 | **** |
|  | Cont vs HSD 2d |  |  | 0.2377 | ns |
|  | Cont vs HSD 5d |  |  | <0.0001 | **** |
|  | Cont vs HSD 7d |  |  | <0.0001 | **** |
|  | HSD+veh 2d vs HSD+veh 5d |  |  | 0.0077 | ** |
|  | HSD+veh 2d vs HSD+veh 7d |  |  | <0.0001 | **** |
|  | HSD+veh 5d vs HSD+veh 7d |  |  | <0.0001 | **** |
| Fig. S2C | Cell number: Cont vs HSD | unpaired 2-tailed t-test |  | 0.209366 | ns |
| Fig. S2C | Cell density: Cont vs HSD | unpaired 2-tailed t-test |  | 0.795484 | ns |
| Fig. S2D | Cell number: Cont vs HSD | unpaired 2-tailed t-test |  | 0.535181 | ns |
| Fig. S2D | Cell density: Cont vs HSD | unpaired 2-tailed t-test |  | 0.157220 | ns |
| Fig. S3B |  | 1-way ANOVA | Tukey's | <0.0001 | **** |
|  | Cont vs HSD 2d |  |  | 0.0589 | ns |
|  | Cont vs HSD 5d |  |  | 0.0404 | * |
|  | Cont vs HSD 7d |  |  | <0.0001 | **** |
|  | HSD+veh 2d vs HSD+veh 5d |  |  | 0.9985 | ns |
|  | HSD+veh 2d vs HSD+veh 7d |  |  | 0.1111 | ns |
|  | HSD+veh 5d vs HSD+veh 7d |  |  | 0.1511 | ns |
| Fig. S3C |  | 1-way ANOVA | Tukey's | <0.0001 | **** |
|  | Cont vs HSD 2d |  |  | 0.9959 | ns |
|  | Cont vs HSD 5d |  |  | 0.7739 | ns |
|  | Cont vs HSD 7d |  |  | <0.0001 | **** |
|  | HSD+veh 2d vs HSD+veh 5d |  |  | 0.9074 | ns |
|  | HSD+veh 2d vs HSD+veh 7d |  |  | 0.0005 | *** |
|  | HSD+veh 5d vs HSD+veh 7d |  |  | 0.0029 | ** |
| Fig. S3D |  | 2-way ANOVA | Tukey's | <0.0001 | **** |
|  | Cont vs HSD 2d |  |  | 0.1947 | ns |
|  | Cont vs HSD 5d |  |  | <0.0001 | **** |
|  | Cont vs HSD 7d |  |  | <0.0001 | **** |
|  | HSD+veh 2d vs HSD+veh 5d |  |  | 0.0058 | ** |
|  | HSD+veh 2d vs HSD+veh 7d |  |  | <0.0001 | **** |
|  | HSD+veh 5d vs HSD+veh 7d |  |  | <0.0001 | **** |
| Fig. S4C | Cell number: Cont vs HSD | unpaired 2-tailed t-test |  | 0.058380 | ns |
| Fig. S4C | Cell density: Cont vs HSD | unpaired 2-tailed t-test |  | 0.785646 | ns |
| Fig. S4D | inside SON vs outside SON | 2-way ANOVA | Šídák's | <0.0001 | **** |
| Fig. S5B |  | 1-way ANOVA | Tukey's | <0.0001 | **** |
|  | Cont+veh vs HSD+veh |  |  | <0.0001 | **** |
|  | Cont+veh vs HSD+PLX3397 |  |  | 0.9218 | ns |
|  | HSD+veh vs HSD+ PLX3397 |  |  | 0.0002 | *** |
| Fig. S5C |  | 1-way ANOVA | Tukey's | 0.0002 | *** |
|  | Cont+veh vs HSD+veh |  |  | 0.0004 | *** |
|  | Cont+veh vs HSD+ PLX3397 |  |  | 0.9174 | ns |
|  | HSD+veh vs HSD+ PLX3397 |  |  | 0.0010 | *** |
| Fig. S5D |  | 1-way ANOVA | Tukey's | <0.0001 | **** |
|  | Cont+veh vs HSD+veh |  |  | <0.0001 | **** |
|  | Cont+veh vs HSD+ PLX3397 |  |  | 0.9776 | ns |
|  | HSD+veh vs HSD+ PLX3397 |  |  | 0.0001 | *** |
| Fig. S6B |  | 1-way ANOVA | Tukey's | <0.0001 | **** |
|  | Cont+veh vs HSD+veh |  |  | 0.0005 | *** |
|  | Cont+veh vs HSD+ PLX5622 |  |  | 0.0045 | ** |
|  | HSD+veh vs HSD+ PLX5622 |  |  | <0.0001 | **** |
| Fig. S6C |  | 1-way ANOVA | Tukey's | 0.0002 | *** |
|  | Cont+veh vs HSD+veh |  |  | 0.0182 | * |
|  | Cont+veh vs HSD+ PLX5622 |  |  | 0.0041 | ** |
|  | HSD+veh vs HSD+ PLX5622 |  |  | <0.0001 | **** |
| Fig. S6D |  | 1-way ANOVA | Tukey's | 0.004 | *** |
|  | Cont+veh vs HSD+veh |  |  | 0.0006 | *** |
|  | Cont+veh vs HSD+ PLX5622 |  |  | 0.7896 | ns |
|  | HSD+veh vs HSD+ PLX5622 |  |  | 0.0015 | ** |
| Fig. S7B |  | 1-way ANOVA | Tukey's | <0.0001 | **** |
|  | Cont+veh vs HSD+veh |  |  | <0.0001 | **** |
|  | Cont+veh vs HSD+ BLZ |  |  | 0.6713 | ns |
|  | HSD+veh vs HSD+ BLZ |  |  | <0.0001 | **** |
| Fig. S7C |  | 1-way ANOVA | Tukey's | <0.0001 | **** |
|  | Cont+veh vs HSD+veh |  |  | <0.0001 | **** |
|  | Cont+veh vs HSD+ BLZ |  |  | 0.0785 | ns |
|  | HSD+veh vs HSD+ BLZ |  |  | <0.0001 | **** |
| Fig. S7D |  | 1-way ANOVA | Tukey's | <0.0001 | **** |
|  | Cont+veh vs HSD+veh |  |  | <0.0001 | **** |
|  | Cont+veh vs HSD+ BLZ |  |  | 0.0164 | * |
|  | HSD+veh vs HSD+ BLZ |  |  | 0.0022 | ** |
| Fig. S8B | 2 days | 1-way ANOVA | Tukey's | 0.8080 | ns |
|  | Cont+veh vs HSD+veh |  |  | 0.7967 | ns |
|  | Cont+veh vs HSD+ BLZ |  |  | 0.9059 | ns |
|  | HSD+veh vs HSD+ BLZ |  |  | 0.9732 | ns |
| Fig. S8B | 5 days | 1-way ANOVA | Tukey's | 0.6780 | ns |
|  | Cont+veh vs HSD+veh |  |  | 0.8763 | ns |
|  | Cont+veh vs HSD+ BLZ |  |  | 0.9323 | ns |
|  | HSD+veh vs HSD+ BLZ |  |  | 0.6547 | ns |
| Fig. S8B | 7 days | 1-way ANOVA | Tukey's | 0.0053 | ** |
|  | Cont+veh vs HSD+veh |  |  | 0.0050 | ** |
|  | Cont+veh vs HSD+ BLZ |  |  | 0.6300 | ns |
|  | HSD+veh vs HSD+ BLZ |  |  | 0.0421 | * |
| Fig. S8C | 2 days | 1-way ANOVA | Tukey's | 0.3678 | ns |
|  | Cont+veh vs HSD+veh |  |  | 0.3889 | ns |
|  | Cont+veh vs HSD+ BLZ |  |  | 0.4957 | ns |
|  | HSD+veh vs HSD+ BLZ |  |  | 0.9789 | ns |
| Fig. S8C | 5 days | 1-way ANOVA | Tukey's | 0.0139 | * |
|  | Cont+veh vs HSD+veh |  |  | 0.0320 | * |
|  | Cont+veh vs HSD+ BLZ |  |  | 0.9763 | ns |
|  | HSD+veh vs HSD+ BLZ |  |  | 0.0212 | * |
| Fig. S8C | 7 days | 1-way ANOVA | Tukey's | 0.0023 | ** |
|  | Cont+veh vs HSD+veh |  |  | 0.0098 | ** |
|  | Cont+veh vs HSD+ BLZ |  |  | 0.7475 | ns |
|  | HSD+veh vs HSD+ BLZ |  |  | 0.0032 | ** |
| Fig. S8D | 2 days | 1-way ANOVA | Tukey's | 0.1430 | ns |
|  | Cont+veh vs HSD+veh |  |  | 0.1482 | ns |
|  | Cont+veh vs HSD+ BLZ |  |  | 0.2800 | ns |
|  | HSD+veh vs HSD+ BLZ |  |  | 0.9151 | ns |
| Fig. S8D | 5 days | 1-way ANOVA | Tukey's | 0.0003 | *** |
|  | Cont+veh vs HSD+veh |  |  | 0.0004 | *** |
|  | Cont+veh vs HSD+ BLZ |  |  | 0.6238 | ns |
|  | HSD+veh vs HSD+ BLZ |  |  | 0.0018 | ** |
| Fig. S9D | 7 days | 1-way ANOVA | Tukey's | <0.0001 | **** |
|  | Cont+veh vs HSD+veh |  |  | <0.0001 | **** |
|  | Cont+veh vs HSD+ BLZ |  |  | 0.0031 | ** |
|  | HSD+veh vs HSD+ BLZ |  |  | <0.0001 | **** |
| Fig. S9A |  | 1-way ANOVA | Tukey's | <0.0001 | **** |
|  | Cont+veh vs HSD+veh |  |  | <0.0001 | **** |
|  | Cont+veh vs HSD+ BLZ |  |  | 0.1062 | ns |
|  | HSD+veh vs HSD+ BLZ |  |  | 0.0003 | *** |
| Fig. S9B |  | 1-way ANOVA | Tukey's | 0.0015 | ** |
|  | Cont+veh vs HSD+veh |  |  | <0.0052 | ** |
|  | Cont+veh vs HSD+ BLZ |  |  | 0.7729 | ns |
|  | HSD+veh vs HSD+ BLZ |  |  | 0.0020 | ** |
| Fig. S9C |  | 1-way ANOVA | Tukey's | 0.0003 | *** |
|  | Cont+veh vs HSD+veh |  |  | 0.0003 | *** |
|  | Cont+veh vs HSD+ BLZ |  |  | 0.3127 | ns |
|  | HSD+veh vs HSD+ BLZ |  |  | 0.0019 | ** |
| Fig. S9D |  | 1-way ANOVA | Tukey's | 0.0003 | *** |
|  | Cont+veh vs HSD+veh |  |  | 0.0006 | *** |
|  | Cont+veh vs HSD+ BLZ |  |  | 0.9987 | ns |
|  | HSD+veh vs HSD+ BLZ |  |  | 0.0005 | *** |
| Fig. S10A | Cont vs HSD | unpaired 2-tailed t-test |  | <0.0001 | **** |
| Fig. S10B | Cont vs HSD | unpaired 2-tailed t-test |  | 0.0587 | ns |
| Fig. S10C | Cont vs HSD | unpaired 2-tailed t-test |  | 0.1169 | ns |
| Fig. S11E |  | 2-way RM ANOVA | Tukey's | <0.0001 | **** |
|  | baseline |  |  |  |  |
|  | Cont+veh vs HSD+veh |  |  | 0.9998 | ns |
|  | Cont+veh vs Cont+BLZ |  |  | 0.9001 | ns |
|  | Cont+veh vs HSD+BLZ |  |  | 0.3287 | ns |
|  | HSD+veh vs Cont+BLZ |  |  | 0.8650 | ns |
|  | HSD+veh vs HSD+BLZ |  |  | 0.2895 | ns |
|  | Cont+BLZ vs HSD+BLZ |  |  | 0.7217 | ns |
|  | DHK |  |  |  |  |
|  | Cont+veh vs HSD+veh |  |  | <0.0001 | **** |
|  | Cont+veh vs Cont+BLZ |  |  | 0.4153 | ns |
|  | Cont+veh vs HSD+BLZ |  |  | 0.9855 | ns |
|  | HSD+veh vs Cont+BLZ |  |  | <0.0001 | **** |
|  | HSD+veh vs HSD+BLZ |  |  | <0.0001 | **** |
|  | Cont+BLZ vs HSD+BLZ |  |  | 0.6174 | ns |
| Fig. S11F |  | 2-way RM ANOVA | Tukey's | <0.0001 | **** |
|  | baseline |  |  |  |  |
|  | Cont+veh vs HSD+veh |  |  | 0.8963 | ns |
|  | Cont+veh vs Cont+BLZ |  |  | 0.4081 | ns |
|  | Cont+veh vs HSD+BLZ |  |  | 0.7652 | ns |
|  | HSD+veh vs Cont+BLZ |  |  | 0.1339 | ns |
|  | HSD+veh vs HSD+BLZ |  |  | 0.3619 | ns |
|  | Cont+BLZ vs HSD+BLZ |  |  | 0.9276 | ns |
|  | DHK |  |  |  |  |
|  | Cont+veh vs HSD+veh |  |  | <0.0001 | **** |
|  | Cont+veh vs Cont+BLZ |  |  | 0.7769 | ns |
|  | Cont+veh vs HSD+BLZ |  |  | 0.9274 | ns |
|  | HSD+veh vs Cont+BLZ |  |  | <0.0001 | **** |
|  | HSD+veh vs HSD+BLZ |  |  | <0.0001 | **** |
|  | Cont+BLZ vs HSD+BLZ |  |  | 0.9858 | ns |
| Fig. S12E |  | 2-way RM ANOVA | Tukey's | <0.0001 | **** |
|  | baseline |  |  |  |  |
|  | Cont+veh vs HSD+veh |  |  | >0.9999 | ns |
|  | Cont+veh vs HSD+BLZ |  |  | 0.5451 | ns |
|  | Cont+veh vs Cont+BLZ |  |  | 0.9992 | ns |
|  | HSD+veh vs HSD+BLZ |  |  | 0.5670 | ns |
|  | HSD+veh vs Cont+BLZ |  |  | 0.9983 | ns |
|  | HSD+BLZ vs Cont+BLZ |  |  | 0.4677 | ns |
|  | ifenprodil |  |  |  |  |
|  | Cont+veh vs HSD+veh |  |  | <0.0001 | **** |
|  | Cont+veh vs HSD+BLZ |  |  | 0.7479 | ns |
|  | Cont+veh vs Cont+BLZ |  |  | 0.9855 | ns |
|  | HSD+veh vs HSD+BLZ |  |  | <0.0001 | **** |
|  | HSD+veh vs Cont+BLZ |  |  | <0.0001 | **** |
|  | HSD+BLZ vs Cont+BLZ |  |  | 0.7508 | ns |
| Fig. S12F |  | 2-way RM ANOVA | Tukey's | <0.0001 | **** |
|  | baseline |  |  |  |  |
|  | Cont+veh vs HSD+veh |  |  | 0.9870 | ns |
|  | Cont+veh vs HSD+BLZ |  |  | 0.1019 | ns |
|  | Cont+veh vs Cont+BLZ |  |  | 0.7681 | ns |
|  | HSD+veh vs HSD+BLZ |  |  | 0.1864 | ns |
|  | HSD+veh vs Cont+BLZ |  |  | 0.9189 | ns |
|  | HSD+BLZ vs Cont+BLZ |  |  | 0.4777 | ns |
|  | ifenprodil |  |  |  |  |
|  | Cont+veh vs HSD+veh |  |  | <0.0001 | **** |
|  | Cont+veh vs HSD+BLZ |  |  | 0.0782 | ns |
|  | Cont+veh vs Cont+BLZ |  |  | 0.9921 | ns |
|  | HSD+veh vs HSD+BLZ |  |  | <0.0001 | **** |
|  | HSD+veh vs Cont+BLZ |  |  | <0.0001 | **** |
|  | HSD+BLZ vs Cont+BLZ |  |  | 0.1333 | ns |

## STAR Methods

### Animals

Adult Wistar rats (300-400 g; Charles River Laboratories Inc., Saint-Constant, Quebec) were used throughout this study. Rats were treated in strict accordance with the guidelines outlined by the Canadian Council on Animal Care (http://www.ccac.ca/) and experiments adhered to protocols #AUP7948 approved by the Facility Animal Care Committee of McGill University. All animals were kept in a climate-controlled room under a standard 12/12h light/dark cycle, habituated to individual housing in behavioral cages for 10 days before experimental procedures (PhenoMaster, TSE Systems). Food and water were provided *ad libitum*. HSD-treated rats were provided with 2% NaCl solution as drinking fluid instead of water for 2-7 days as described previously^38,39,79,80^. Electrophysiological experiments were performed on in-house bred transgenic Wistar rats (8-12 weeks old) expressing enhanced green fluorescent protein (eGFP) under the control of the Avp gene promoter^45^. In all experiments, animals were randomly assigned to different groups. The experimenter was blinded to the experimental condition in all studies.

### Immunohistochemistry on tissue sections

Rats were transcardially perfused with 10 ml of phosphate-buffered saline (PBS) and then with 250 ml of PBS containing 4% paraformaldehyde and 0.25% glutaraldehyde, pH adjusted to 6.8, at 37°C. Brains were post-fixed for at least 48 hours. Coronal sections (50 μm thick) were obtained using a vibratome and were treated for 15 min with 0.1% NaBH_4_ in cytoskeletal buffer containing (in mM): NaCl 130, MES 10, EGTA 5, MgCl_2_ 5, Glucose 5, pH adjusted to 6.3. After one-hour incubation with 10% normal goat serum in PBS containing 0.3% Triton-X, sections were washed and incubated for 24 hours at 4°C with primary antibodies diluted in PBS containing 0.3% Triton-X. For incubation with primary antibody against GLT1, 5% normal goat serum was added to the primary antibody solution. Following washes with PBS, sections were incubated for two hours with fluorescently labelled secondary antibodies, mounted in ProLong Gold Antifade reagent (Life Technologies), and imaged using LSM 880 with AiryScan (Zeiss).

The images presented in the figures illustrate the middle parts of the SON and the magnocellular PVN, at anterior-posterior bregma coordinates -0.90 to -1.00 mm for the SON, and -1.70 to -1.80 mm for the PVN. The morphology of the SON and PVN appears different, specifically, both nuclei are bigger with more prominent VP neuron population and angiogenesis due to the previously characterized phenomenon referred to as hypertrophy, that takes place in these nuclei in response _to HSD8,81,82._

### Antibodies

Primary antibodies used were vasopressin guinea pig polyclonal antibody (SYSY 403004, 1:500), oxytocin mouse monoclonal antibody (PS38, 1:100; provided by Dr. Hal Gainer, NIH), GFAP chicken polyclonal antibody (Abcam ab4674, 1:1000), Iba1 rabbit polyclonal antibody (WAKO 019-19741, 1:500), Iba1 guinea pig polyclonal antibody (SYSY 234-004, 1:500), CD68 mouse monoclonal antibody (Biorad MCA341GA, 1:500), S100β rabbit monoclonal antibody (Abcam ab52642, 1:500), ALDH1L1 rabbit polyclonal antibody (Proteintech 17390-1-AP, 1:50), GLT1 rabbit polyclonal antibody (Abcam ab106289, 1:500), vimentin mouse monoclonal antibody (DSHB 40ECs, 1:100), TMEM119 rabbit polyclonal antibody (Synaptic Systems 400203, 1:400), NeuN guinea pig polyclonal antibody (Millipore Sigma ABN90, 1:200). Secondary antibodies used were Alexa-conjugated IgG (405 nm, 488 nm, 568 nm, and 647 nm; 1:500, Invitrogen) and DAPI (1:5000, Invitrogen).

### Confocal microscopy with AiryScan

Confocal microscopy was conducted using LSM 880 with Apo 20x/0.8 objective (Zeiss), and AiryScan imaging was carried out using Zeiss 63x/1.40 Oil DIC f/ELYRA objective. For the super-resolution imaging, the AiryScan super-resolution (SR) module with 32-channel hexagonal array GaAsP detector for LSM (Zeiss) was used to collect stacks of 100-150 optical sections (170-nm step) using imaging parameters as we previously described^80^. The point spread functions (PSFs) were measured and the maximal resolution of the Airyscan system at the wavelength used for CD68 imaging (excitation of 488 nm) was 126 nm in the lateral (xy) dimension and 346 nm in the axial (z) dimension. AiryScan super-resolution image stacks were reconstructed using ZEN Black software (Zeiss). Images were analyzed using ZEN Blue software (Zeiss) and FIJI (NIH).

### Image analysis and quantification

For the glia coverage analysis, 50 μm-thick brain sections containing SON or PVN were imaged as z-stacks (1 μm step, 20x objective). Maximal intensity projection images were generated and the SON or PVN area containing VP neurons was identified by outlining the location of VP-positive cells. Then, the total immunopositive area for Iba1 or GFAP overlapping with the location of VP neurons was calculated using FIJI. Microglia density was calculated by counting the number of Iba1-positive cells in a 100 μm x 100 μm area positioned over a central region of VP-positive area in the SON or PVN.

The intensity of S100β, ADLH1L1, GLT1, and vimentin in the SON is presented as a percentage relative to the area outside the SON. The relative intensity was calculated by dividing the total intensity of the SON VP-positive area by the total intensity of the adjacent VP-negative area outside the SON.

The total number of astrocytes in the SON and PVN was calculated by counting the number of S100β-positive cells within the area outlined by the location of VP-positive cells in the SON or the magnocellular part of the PVN. Astrocyte density in the SON and PVN was calculated by dividing the total number of S100β-positive cells by the VP-positive corresponding area.

For analysing microglia morphology, brain sections labeled for the VP, Iba1, and DAPI, z-stacks (60-100 optical sections, 0.5 μm step, 63x objective) were imaged. Microglia shape was analysed in animals from different experimental groups, comparing morphology of Iba1-positive cells located within the area containing VP neurons and outside the SON and PVN. The microglia complexity and process ramification were evaluated by calculating the number of Sholl intersections as a function of the distance from the microglial soma. For the generation and comparison of Sholl distribution curves, individual microglia Sholl profiles were generated, and profiles of 20 cells were averaged per rat^83^. Each data point represents a mean value calculated for 5 rats per condition (Fig. 2d). Incompletely stained microglia were not included in the analysis.

For the CD68 lysosome quantification, z-stacks containing the whole microglia soma were acquired using AiryScan super-resolution mode (150 nm step, 63x objective, zoom 2x). Maximal projection images were generated, and the area occupied by CD68 positive lysosomes was calculated for an individual microglia soma as a ratio between total CD68+ area and the total Iba1-positive area. FIJI 3D reconstruction of Iba1-positive microglia was used to identify and quantify the location of astrocytic and neuronal markers inside phagocytic lysosomes. Then, the number of CD68+ lysosomes, CD68/GFAP-double positive lysosomes, CD68/S100β-double positive lysosomes, CD68/NeuN-double positive lysosomes, and CD68/VP-double positive lysosomes were counted for individual microglia.

IMARIS 3D Reconstruction of microglia were done by converting raw czi stacks of to the ims format in IMARIS software (Version 10.1.1, Oxford Instruments). Image processing was used to remove background noise prior to reconstruction by applying baseline subtraction to the Iba1 channel. Microglia and lysosomes were reconstructed using the surfaces tool by using the smooth function followed by absolute intensity thresholding. For astrocytic proteins (GFAP and S100β), the thresholding was performed using background subtraction (local contrast). Videos were made in the Animation window and images of the reconstructed microglia were taken using the Snapshot tool.

### Administration of microglia inhibitors

PLX3397 (MedChemExpress) was first diluted as a stock in dimethyl sulfoxide (DMSO, 100 mg/ml), sonicated, aliquoted, and stored at -80 °C. Prior to the drug administration, the stock was diluted to prepare the injecting solution in the final concentration of 10 mg/ml in 10% DMSO, 40% PEG300, 5% Tween-80, and 45% saline. Rats (∼300-350 g) were i.p. administered with 1 ml of the injection solution daily, for the final dose of 30 mg/kg/day. The control group received 1 ml of the vehicle containing 10% DMSO, 40% PEG300, 5% Tween-80, and 45% saline.

PLX5622 (Cat.#HY-114153, MedChemExpress) was first diluted as a stock in dimethyl sulfoxide (DMSO, 100 mg/ml), sonicated, aliquoted, and stored at -80 °C. Prior to the drug administration, the stock was diluted to prepare the injecting solution in the final concentration of 6.5 mg/ml in 10% DMSO, 20% RH40 (KolliphorR RH40, Sigma), 5% Tween-80 and 65% saline. Rats (∼300-350 gr) were i.p. administered with 2.7 ml of the injection solution twice a day, for the final dose of 100 mg/kg/day. The control group received 2.7 ml of the vehicle containing 10% DMSO, 20% RH40, 5% Tween-80, and 65% saline.

BLZ945 (MedChemExpress) was dissolved as a stock in DMSO at the concentration of 60 mg/ml and stored at -80 °C. Then, 2 μl of the stock solution was mixed with 4 μl of PEG300, and the total 6 μl solution (final concentration of 20 mg/ml) was injected i.c.v. into rats daily. The control group received 6 µl of the vehicle containing 2 µl DMSO and 4 µl PEG300 daily. Rats were installed with i.c.v. cannulas ∼ 2 weeks prior to the first BLZ945 or vehicle injection, and i.c.v. injections were done in awake freely moving rats.

All the drugs or vehicles were administered for 3 days prior to introducing rats to HSD or control diet, and drug or vehicle administration continued during the entire period when rats were exposed to control or HSD. Both i.p. and i.c.v. drug and vehicle administrations were performed in awake rats. To habituate them to the procedure, the rats were handled for 5 days prior to the first manipulation.

### Acute hypothalamic slice preparations

VP-eGFP rats were euthanized by decapitation and brains were isolated and kept in an ice-cold sucrose solution containing (in mM): 252 sucrose, 2.5 KCl, 2 MgCl_2_, 1.5 CaCl_2_, 1.25 NaH_2_PO_4_, 26 NaHCO_3_, and 10 D-glucose; pH 7.35; 300–320 mOsmol/kg, bubbled with carbogen (95% O_2_ and 5% CO_2_). Acute angled horizontal slices (400 μm thick) were prepared as described previously^84^ and placed in a recording chamber on a fixed-stage microscope equipped with a 40X water immersion objective (Zeiss examiner, Z1. AXIO, Zeiss). Slices were perfused at 1.5 ml/min with warm oxygenated (95% O_2_; 5% CO_2_; 31.5 ± 1°C) artificial cerebrospinal fluid (ACSF) for 45 - 60 minutes before recording. The ACSF comprised (in mM) 104 NaCl, 26 NaHCO_3_, 1.2 NaH_2_PO_4_, 3 KCl, 1 MgCl_2_, 2 CaCl_2_, 10 D-glucose, osmolality 297 ± 3 mOsmol/kg.

### Whole-cell electrophysiology

For current clamp recordings to measure the neuronal excitability, the patch pipettes were filled with intracellular solution containing (in mM): 140 K-gluconate, 2.5 MgCl_2_, 10 HEPES, 2 Na_2_-ATP, 0.5 Na-GTP, and 0.5 EGTA; pH 7.3, 285–295 mOsm/kg. Pipette resistance in the bath was 3–4 MΩ and recordings were excluded if the RMP was more positive than −50 mV or series resistance was >25 MΩ. Baseline activity was recorded in ACSF for at least 15 minutes before a drug application of glutamate transporter 1 (GLT1) inhibitor dihydrokainic acid (DHK 100 μM, Sigma) or NMDAR subunit NR2B antagonist ifenprodil (10 μM, Sigma).

For voltage clamp recordings to measure the NMDA receptor-mediated tonic current, the patch pipettes were filled with an intracellular solution containing (in mM): 135 cesium methane-sulphonate, 0.2 EGTA, 10 HEPES, 1 MgCl_2_, 2 MgATP, 0.2 NaGTP, 10 TEA-Cl, and 20 sodium phosphocreatine; pH 7.25 ± 0.05. The low magnesium ACSF used in recording NMDA currents comprised (in mM): 104 NaCl, 26 NaHCO_3_, 1.2 NaH_2_PO_4_, 3 KCl, 0.02 MgCl_2_, 2 CaCl_2_, 10 D-glucose, and 0.001 tetrodotoxin (TTX, Alomone labs, Israel), osmolality 297 ± 3 mOsmol/kg.

The NMDA receptor-mediated tonic current (tonic I_NMDA_) was assessed as changes in the holding current (I_holding_). The mean tonic I_NMDA_ amplitude was calculated by the difference in I_holding_ measured as the average of a 2 min steady-state baseline segment obtained before and 5 min after the application of NMDAR antagonists (ifenprodil 10 μM, AP-5 100 μM, and kynurenic acid 1mM, Sigma). I_NMDA_ was recorded and calculated at voltage clamped potential +40 mV.

The data were acquired with pCLAMP 11.0 (Molecular Devices) at a sampling rate of 10-20 kHz and measured and plotted with pCLAMP 11.0. A custom written Matlab script was used for to generate action potential raster plots. Prism (GraphPad Prism 9.1.0) was used to generate histogram distributions.

### Blood pressure monitoring

Blood pressure was monitored by recording arterial blood pressure and heart rate in conscious unrestrained adult rats using radio telemetry as previously described^38^. Adult Wistar rats were anesthetized with isoflurane (induction: 4%, maintenance: 1–2%), and administered with Buphrenorphine (0.05 mg/kg), VetPen 300 (penicillin 1×10^5^ IU/kg), and saline (50 ml/kg). The telemetry transmitter (PTA-M/F FLAT, Stellar Telemetry, TSE Systems) was placed and secured in a subcutaneous pocket on the back of the animal. The attached indwelling catheter with the blood pressure sensor was inserted into the femoral artery and advanced into the abdominal aorta. The systolic and diastolic arterial pressure, heart rate, and body temperature were sampled at 100 Hz for 15-second periods every 30 min. The telemetry sensors wirelessly communicated with an external antenna (Stellar Telemetry, TSE Systems), and the transmitted parameters were recorded by AcqKnowledge BIOPAC software. For the i.c.v. drug administration, 2 weeks after recovery from telemetry implantation surgery, rats were subjected to intracerebroventricular cannula implantations. Blood pressure measurements were initiated a week before treatment onset, to establish baseline arterial pressure values. At the end of dietary and pharmacological protocols, telemetry devices were extracted, and rat brains were fixed to confirm the position of the cannula and evaluate the effects of the treatments.

### Western Blot

Tissue blocks containing the SON were removed and homogenized in ice-cold lysis buffer consisting of 200 mM HEPES, 50 mM NaCl, 10% glycerol, 1% triton X-100, 1 mM EDTA, 50 mM NaE, 2 mM Na_3_VO_4_, 25 mM β-glycerophosphate, and 1 EDTA-free complete ULTRA tablet. A total of 40 μg of protein per sample was resolved on SDS-PAGE (10%), transferred onto nitrocellulose membranes, and imaged using a ChemiDoc Imaging System (BioRad). Primary antibodies for western blotting are GFAP (cat# G4546; Sigma), GLT1 (cat# ab106289; Abcam), ALDH1L1 (cat# 17390-1-AP, Proteintech), raised in rabbit, and β-actin (cat# a5441; Sigma) raised in mouse. Secondary antibody for western blotting is anti-rabbit and anti-mouse HRP (GE Healthcare).

### Statistical analysis

All analyses were performed using Prism (GraphPad 9.1.0). Results are reported as mean plus or minus standard error of the mean (± SEM). Data were assessed for normality and statistical tests were performed to compare group values using Prism. Data of two groups were compared by a two-tailed, unpaired, parametric Student’s t-test. For parametric multiple comparisons, we used one-way ANOVA followed by between-group comparisons using Tukey’s post hoc test, or a repetitive measured (RM) two-way ANOVA followed by between-group multiple comparison Tukey’s or Šídák’s multiple comparisons post-hoc test. Differences between values considered statistically significant at p < 0.05; *p < 0.05, **p < 0.01, ***p < 0.001, ****p < 0.0001. All the analyses, including statistical tests, multiple comparison post-hoc tests (if applicable), and p values for all groups compared in multiple comparisons, are summarised in Table S2.

